# Mutation-Guided Vaccine Design: A Strategy for Developing Boosting Immunogens for HIV Broadly Neutralizing Antibody Induction

**DOI:** 10.1101/2022.11.11.516143

**Authors:** Kevin Wiehe, Kevin O. Saunders, Victoria Stalls, Derek W. Cain, Sravani Venkatayogi, Joshua S. Martin Beem, Madison Berry, Tyler Evangelous, Rory Henderson, Bhavna Hora, Shi-Mao Xia, Chuancang Jiang, Amanda Newman, Cindy Bowman, Xiaozhi Lu, Mary E. Bryan, Joena Bal, Aja Sanzone, Haiyan Chen, Amanda Eaton, Mark A. Tomai, Christopher B. Fox, Ying Tam, Christopher Barbosa, Mattia Bonsignori, Hiromi Muramatsu, S. Munir Alam, David Montefiori, Wilton B. Williams, Norbert Pardi, Ming Tian, Drew Weissman, Frederick W. Alt, Priyamvada Acharya, Barton F. Haynes

## Abstract

A major goal of HIV-1 vaccine development is induction of broadly neutralizing antibodies (bnAbs). While success has been achieved in initiating bnAb B cell lineages, design of boosting immunogens that select for bnAb B cell receptors with improbable mutations required for bnAb affinity maturation remains difficult. Here we demonstrate a process for designing boosting immunogens for a V3-glycan bnAb B cell lineage. The immunogens induced affinity-matured antibodies by selecting for functional improbable mutations in bnAb precursor knock-in mice. Moreover, we show similar success in prime and boosting with nucleoside-modified mRNA-encoded HIV-1 envelope trimer immunogens, with improved selection by mRNA immunogens of improbable mutations required for bnAb binding to key envelope glycans. These results demonstrate the ability of both protein and mRNA prime-boost immunogens for selection of rare B cell lineage intermediates with neutralizing breadth after bnAb precursor expansion, a key proof-of concept and milestone towards development of an HIV vaccine.

**One-Sentence Summary:** A vaccine strategy for selecting key rare antibody mutations is shown to induce HIV broadly neutralizing antibodies in mice.

## Main Text

Induction of broadly neutralizing antibodies (bnAbs) has emerged as the primary goal of HIV-1 vaccine development efforts (*1*, *2*). BnAb induction can occur in HIV-1 infection, but only after a series of rare events occur, including evolution of envelope (Env) variants that can bind to naïve and early lineage bnAb B cells, expansion of rare precursors, and selection of bnAb B cell receptors (BCRs) with improbable mutations that are required for affinity maturation to full bnAb breadth and potency (*3*–*5*). Improbable mutations are rarely made by activation-induced cytidine deaminase during affinity maturation but can result in amino acid substitutions that improve antibody affinity for antigen and confer broad neutralization activity (*4*–*6*). To convert a series of rare events to become more common with vaccination, a strategy has been proposed to use a series of sequential immunogens designed to bind to bnAb B cell lineage naïve B cell precursors and to select intermediate BCRs with the desired improbable functional mutations (*3–5*). Success in this endeavor requires design of boosting immunogens that select for bnAb lineage intermediate antibodies with mutations required for bnAb B cell receptor interactions with both proteins and glycans.

Computationally reconstructed trees representing B cell clonal genealogies have been central to the study of antibody evolution and have informed HIV-1 vaccine design strategies for induction of bnAbs (*2*, *3*, *5*, *7*–*19*). The inferred unmutated common ancestor (UCA) at the root of a bnAb clonal tree is a representation of the original B cell progenitor of the bnAb B cell clone, and recent successes in designing immunogens that can engage B cell germlines and expand bnAb precursor B cells have been reported (*6*, *11*, *20*–*26*). However, a major roadblock following germline targeting of bnAb precursor B cells is lack of a strategy to design boosting immunogens to select for required improbable functional mutations (*2*, *6*, *20*).

The DH270 bnAb B cell clone was isolated from an HIV-1-infected subject after acute HIV-1 infection and targets the conserved V3 glycan bnAb epitope (*5*). We designed a priming immunogen (10.17DT) for optimal DH270 UCA binding by removing two key glycans from the first variable (V1) loop of the HIV-1 Env (10.17DT) (*6*). Here we use the DH270 UCA V_H_^(+,-)^/V_L_^(+,-)^ knock-in mouse model to show the process for design and interrogation of boosting immunogens following a germline-targeting prime needed to select for accumulation of multiple bnAb functional improbable mutations. Moreover, we show that both protein and nucleoside-modified mRNA-encoded Env boosts selected for functional improbable mutations in bnAb B cell lineage intermediate antibodies. Finally, we show that nucleoside modified mRNA immunization was superior to protein immunization in selecting for BCR improbable mutations required for direct bnAb interactions with Env glycans.

## RESULTS

### Identification of key improbable mutations in DH270 clonal lineage intermediates

To identify where improbable mutation bottlenecks occurred in the maturation of the DH270 bnAb B cell clonal lineage, we mapped the probability of each mutation along the maturation pathway from the DH270 UCA to the broadest and most potent member of the DH270 clone (DH270.6) using ARMADiLLO, a computational program that estimates the probability of amino acid substitutions by simulating somatic hypermutation in the absence of selection (*4*).The maturation pathway includes 46 amino acid changes, of which 22 were classified by ARMADiLLO as improbable (<2% probability of occurring in the absence of selection) (**Figure 1A**). We have recently shown that many of the mutations that occurred in DH270.6 are not required for heterologous neutralization breadth through the design of a minimally mutated DH270 (DH270.min11) which recapitulated 90% of the neutralization capacity of DH270.6 with just 12 mutations (*27*). By mapping the mutation probabilities onto the reconstructed DH270 clonal tree, we determined that half of the minDH270 mutations occurred by the second intermediate (I3.6) in the DH270.6 maturation pathway and all 12 occurred by the third intermediate (I2.6), indicating that acquisition of the most critical mutations occurred during early and mid-stage maturation of the DH270 clone. Additionally, 10 out of 12 critical functional DH270 mutations were improbable within the intermediate antibody in which they first arose, underscoring the importance of the selection of rare mutational events for DH270.6 bnAb development.

**Figure 1.**
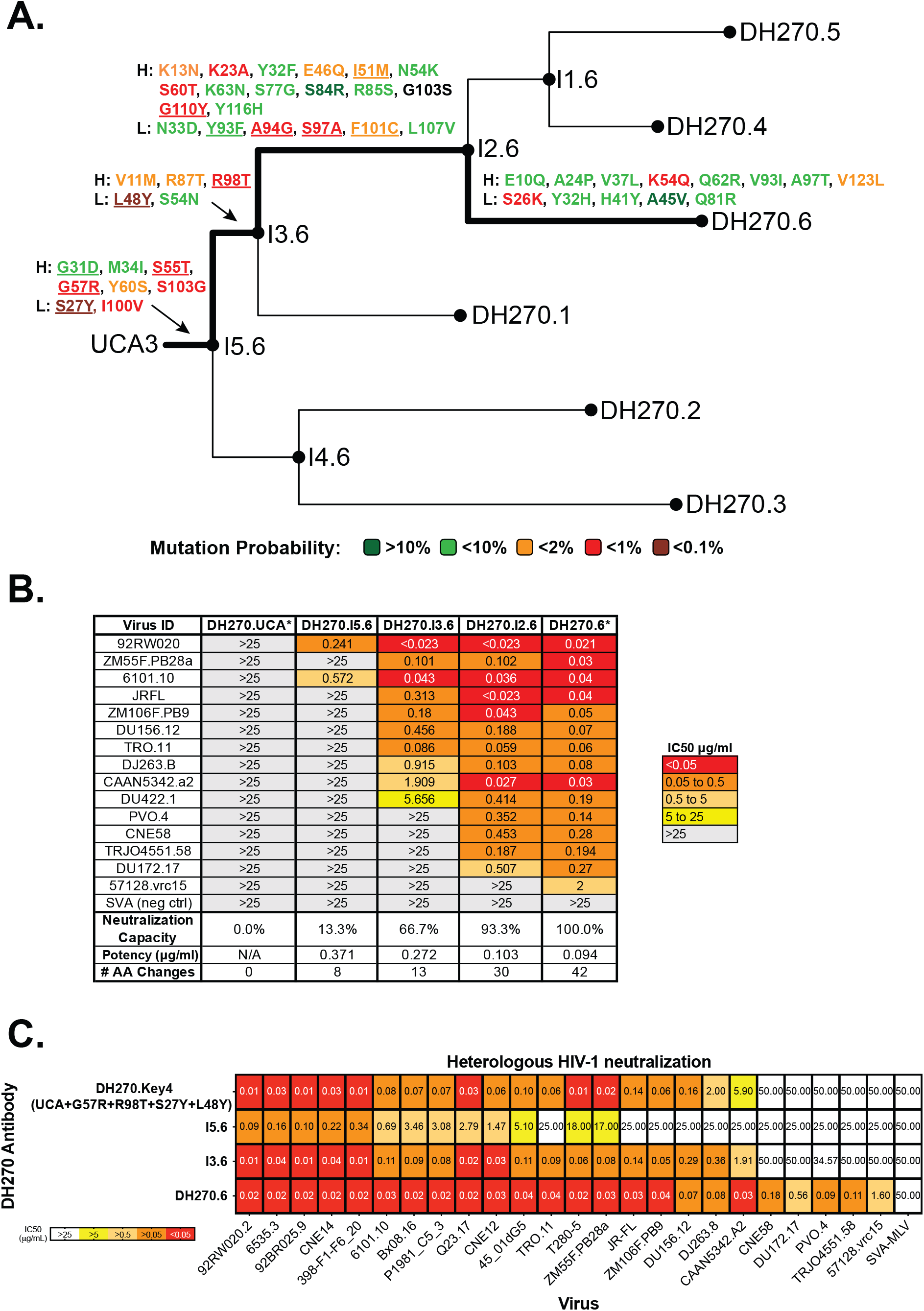
Development of Neutralization Breadth in the DH270.6 Clonal Lineage. **A**) DH270 clonal family tree representing the reconstructed maturation history of the 6 member DH270 clone. Each branch of the maturation pathway from UCA to DH270.6 (bold) is annotated with the individual amino acid changes that occur between the intermediates. Each mutation is colored according to its estimated probability of occurring at that point in the clonal tree in the absence of antigenic selection. The 12 mutations that occur in a minimally mutated DH270 antibody with similar neutralization capacity as DH270.6 (DH270.min11) are underlined. **B)** Neutralization of each ancestral intermediate antibody in the DH270.6 maturation pathway against a panel of viruses selected for sensitivity to DH270.6 neutralization. The neutralization capacity (defined as the percentage of DH270.6-sensitive isolates neutralized by the clonal ancestor), neutralization potency (geometric mean IC50 of detected viruses), and the number of amino acid mutations in each ancestor are listed in the bottom rows. Asterisks denotes that DH270.UCA and DH270.6 neutralization values were originally reported in Bonsignori et al. (*5*). **C)** Introduction of four key improbable mutations (G57R and R98T in heavy chain; S27Y and L48Y in light chain) that occur by the second intermediate (I3.6) into the UCA antibody results in a neutralization profile against an extended viral panel that recapitulates the neutralization profile of the second intermediate.

To determine the critical events in the development of broad neutralization during DH270 clonal maturation, we produced antibodies representing intermediate ancestors along the DH270.6 maturation pathway between the UCA and DH270.6. Neutralization capacity was determined against a 15-virus indicator panel comprised of heterologous viruses sensitive to DH270.6 bnAb neutralization (**Figure 1B**). In contrast to many other bnAb lineages (*13*–*16*, *19*), potent heterologous neutralization in the DH270 V3-glycan B cell lineage was achieved by the first intermediate antibody (*5*). The second intermediate antibody with only 13 mutations had 67% of the neutralization breadth of the DH270.6 bnAb, demonstrating that the largest gain in neutralization capacity during clonal maturation occurred between the first and second intermediates in the DH270.6 lineage (**Figure 1B**).

We asked which of the mutations made the largest contributions to neutralization breath by the second intermediate antibody. There are 5 mutations from the first intermediate antibody to the second intermediate antibody. We introduced every permutation of the 5 mutations into I5.6 (a total of 30 combinations), produced each mutant antibody, and tested their neutralization against viruses sensitive to I3.6 (**Figure S1**). When introduced into I5.6, the combination of two mutations, R98T in the heavy chain and L48Y in the light chain, consistently led to increased neutralization breath. Moreover, reversion of S27Y and L48Y in I3.6 led to the greatest reductions in neutralization breadth (**Figure S1**). Thus, the combined acquisition of R98T and L48Y were the critical mutational events driving the early gain in neutralization breadth in the second intermediate antibody. (**Figure S1**).

We have previously shown that improbable mutations heavy chain G57R and light chain S27Y were also critical for the early heterologous neutralization capacity of the first intermediate (*5*, *6*). Here, we hypothesized that the combined effects of R98T and L48Y with G57R and S27Y would recapitulate the neutralization breadth of the second intermediate. We produced an antibody that introduced these four mutations into the UCA (referred to as DH270.key4) and compared its neutralization capacity with the second intermediate antibody, I3.6. DH270.key4 showed a strikingly similar neutralization pattern to I3.6 on a 24-virus panel of DH270.6-sensitive viruses (**Figure 1C**). Thus, we have identified four key early mutations acquired in combination that can lead to broad and potent heterologous tier 2 virus neutralization. Because all four of the DH270 key mutations are improbable, they are unlikely to occur in combination in a BCR in the germinal center (GC) by chance mutational events. Rather, the appearance of these four mutations in the same BCR would be indicative of strong antigenic selection and on-track evolution of a bnAb B cell clone. Here, we use this set of mutations as markers for monitoring early DH270.6 lineage maturation to assess the effectiveness of vaccines at guiding affinity maturation towards bnAb development.

### Acquisition kinetics of key mutations during immunization

The high mutation frequencies typically observed in bnAbs are likely a byproduct of the extended maturation time required to acquire a restricted set of functional improbable mutations (*4*). Recruitment of memory B cells into secondary GCs is an inefficient process (*28*–*30*) and it has been suggested that sequential vaccine strategies that rely on recall of memory B cells may not be effective for eliciting high levels of SHM (*30*). We and others have hypothesized that an alternative approach to inducing B cells with high mutation levels is through repetitive immunizations with short time intervals to continually stimulate persistent germinal centers thereby prolonging the accumulation of mutations in GC B cells (*6*, *30*–*32*). A frequent, repetitive immunization strategy using 6 immunizations of our 10.17DT nanoparticle was effective at activating DH270 precursor B cells and selecting some early key improbable mutations in DH270 UCA knock-in mice (*6*). To determine if key improbable mutation levels could be attained with fewer than 6 immunizations, we immunized groups of DH270 UCA V_H_^(+,-)^/V_L_^(+,-)^ knock-in mice every 2 weeks with the CH848 10.17DT nanoparticle. Each group received either 1, 2, 3, 4, 5, or 6 immunizations with spleens and draining or systemic lymph nodes (inguinal and axillary, pooled) collected 1 week after the last immunization (**Figure 2A**). We observed an increased frequency of GL7+CD95+ germinal center B cells after the first immunization that reached a plateau after the second immunization (**Figure 2B and S2A**).

**Figure 2.**
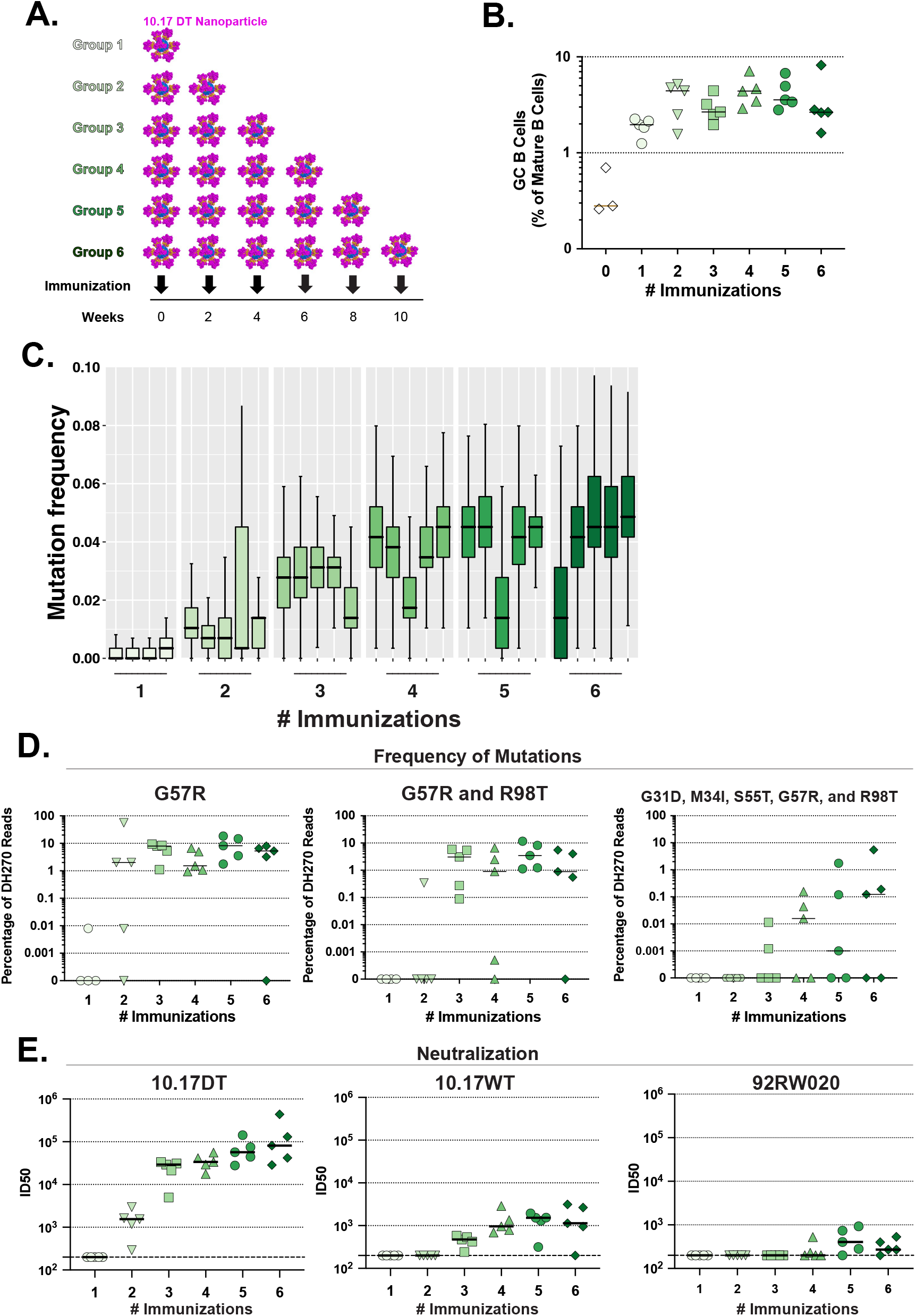
Increasing Numbers of Immunizations Results in Plateaued Response for Repetitive Priming Regimen. **A)** Schematic of immunization regimen in which six groups of DH270 UCA KI mice (n=6 mice per group) were administered 1-6 immunizations of the 10.17DT sortase-conjugated nanoparticles. For each group, mouse BCR repertoires were sequenced 7 days after the last immunization. **B)** Percentage of germinal center B cells in the mature B cell population at 1 week post-immunization increases with the number of immunizations but begins to plateau after the second immunization (Right). Black arrows denote number of immunizations in each mouse group. **C)** Distribution of mutation frequencies of DH270 UCA derived reads from bulk NGS sequencing of heavy chain repertoires of immunized mice. Each box and whiskers plot represents one immunized mouse. Mice are grouped by number of 10.17DT nanoparticle immunizations administered. Mice without sufficient NGS repertoire read counts (<1000 DH270 UCA derived reads) were excluded from analysis (see Methods). **D)** Percentage of DH270 UCA knock-in derived reads in immunized mouse repertoires that possess the heavy chain mutations: improbable G57R (left), co-occurrence of improbable G57R and R98T (middle), and the combination of all heavy chain mutations occurring in the first intermediate of the DH270 clone (right). **E)** Neutralization titers of the vaccine-matched tier 2 10.17DT (left), V1 glycan-containing 10.17 WT (middle) and heterologous tier 1B 92RW020 (right) viruses. Each dot is representative of one immunized DH270 UCA knock-in mouse in panels A, C and D. Bars denote group medians.

Next, we performed bulk unpaired heavy and light chain NGS sequencing of mouse splenocytes to profile the BCR repertoires and DH270 B cell clone mutations. Heavy chain mutation frequency was increased over the first 3 immunizations, but then plateaued (**Figure 2C**). Frequencies of key heavy chain mutations followed a similar pattern, with G57R and R98T frequencies rising and then stabilizing after 3 immunizations both individually and together in combination (**Figure 2D**). Interestingly, the combination of all heavy chain mutations in the first intermediate (I5.6), which includes non-essential DH270 lineage maturation mutations, did increase over the 6 immunizations (**Figure 2D**). For key light chain improbable mutations S27Y and L48Y, the frequency of each increased over the first 3 to 4 immunizations and then plateaued, however the combination of S27Y and L48Y occurring together on the same light chain was only observed in one mouse (**Figure S2B-D**). Mouse serum neutralization of the vaccine-matched virus 10.17DT and the related V1-glycan containing 10.17 WT virus also followed similar kinetics as the mutation frequencies, with titers increasing and then plateauing after 3 immunizations. Heterologous low titer tier 1B 92RW020 virus neutralization was observed in mouse groups that received 5 and 6 immunizations (**Figure 2E**). BCR repertoire sequencing of draining lymph nodes from each mouse in all groups demonstrated comparable mutation patterns and acquisition kinetics in draining lymph node and splenic B cells indicating that, in the setting of repetitive immunizations, mouse splenic B cell repertoire sequencing is sufficiently representative of mutation patterns in draining lymph nodes (**Figure S2E)**.

To determine if these same mutation kinetics occurred in other bnAb B cell lineages, we performed the same studies with the CD4 binding site bnAb CH235 UCA V_H_^(+,-)^/V_L_^(+,-)^ knock-in mouse model (*6*), immunizing 6 groups of mice 1-6 times with the CH505 M5/G458Y/GNTi-germline-targeting SOSIP Env trimer (*6*) and observed a similar pattern of plateauing mutation levels after the 3^rd^ immunization including with the acquisition of a key improbable mutation, K19T (**Figure S2F and G**). CH235 UCA knock-in mouse serum autologous tier 2 neutralization titers increased up to the 4^th^ immunization and then stabilized, a time course similar to that seen in DH270 UCA knock-in mice (**Figure S2H**). These data suggested that for individual or small combinations of key improbable mutations, frequent repetitive immunizations with homologous priming immunogens in bnAb UCA knock-in mice initiated targeted clonal B cell lineages and resulted in rapid gains in key mutation frequencies after 3 immunizations.

### Repetitive immunizations are superior to a single immunization

The frequencies of key mutations we observed in the bnAb UCA knock-in mice may have been influenced not only by the number of immunizations but also by the maturation time from the first immunization. To determine whether time after vaccination also played a role in development of increased frequencies of key mutations, we immunized DH270 UCA knock-in mice once with the 10.17DT nanoparticle and sequenced the BCR repertoires of animals 7 days, 21 days, or 49 days after immunization (**Figure 3A**). For the mouse groups with extended maturation time periods (21 and 49 days) prior to BCR repertoire sequencing, we observed modestly higher overall heavy chain mutation frequencies by 49 days when compared to the mouse group sequenced 7 days post immunization (**Figure 3B**). However, we did not observe any significant differences in the frequencies of total heavy chain improbable mutations or key early lineage heavy chain improbable mutations (**Figures 3C and S3**). In contrast, we observed significantly higher heavy chain mutation frequencies and higher frequencies of key improbable heavy chain mutations in the mouse group immunized 3 times and sequenced 7 days later compared to single immunization mouse groups (p<0.05; Wilcoxon-Mann Whitney exact test) (**Figures 3B and C, and S3**). Moreover, mice immunized 3 times with nanoparticles and sequenced 21 days later also did not result in higher mutation levels relative to mice immunized 3 times and sequenced 7 days later, again indicating that time duration was not the main driver of increased somatic mutation in GC B cells (**Figures 3B and C, and S3**). We conclude that continual antigenic fueling of GCs by frequent repetitive immunizations induced B cells with high mutation frequencies, as well as increased levels of SHM in DH270 UCA knock-in mice were primarily influenced by the number of immunizations over a short period of time and were less a product of prolonged maturation time after immunization.

**Figure 3.**
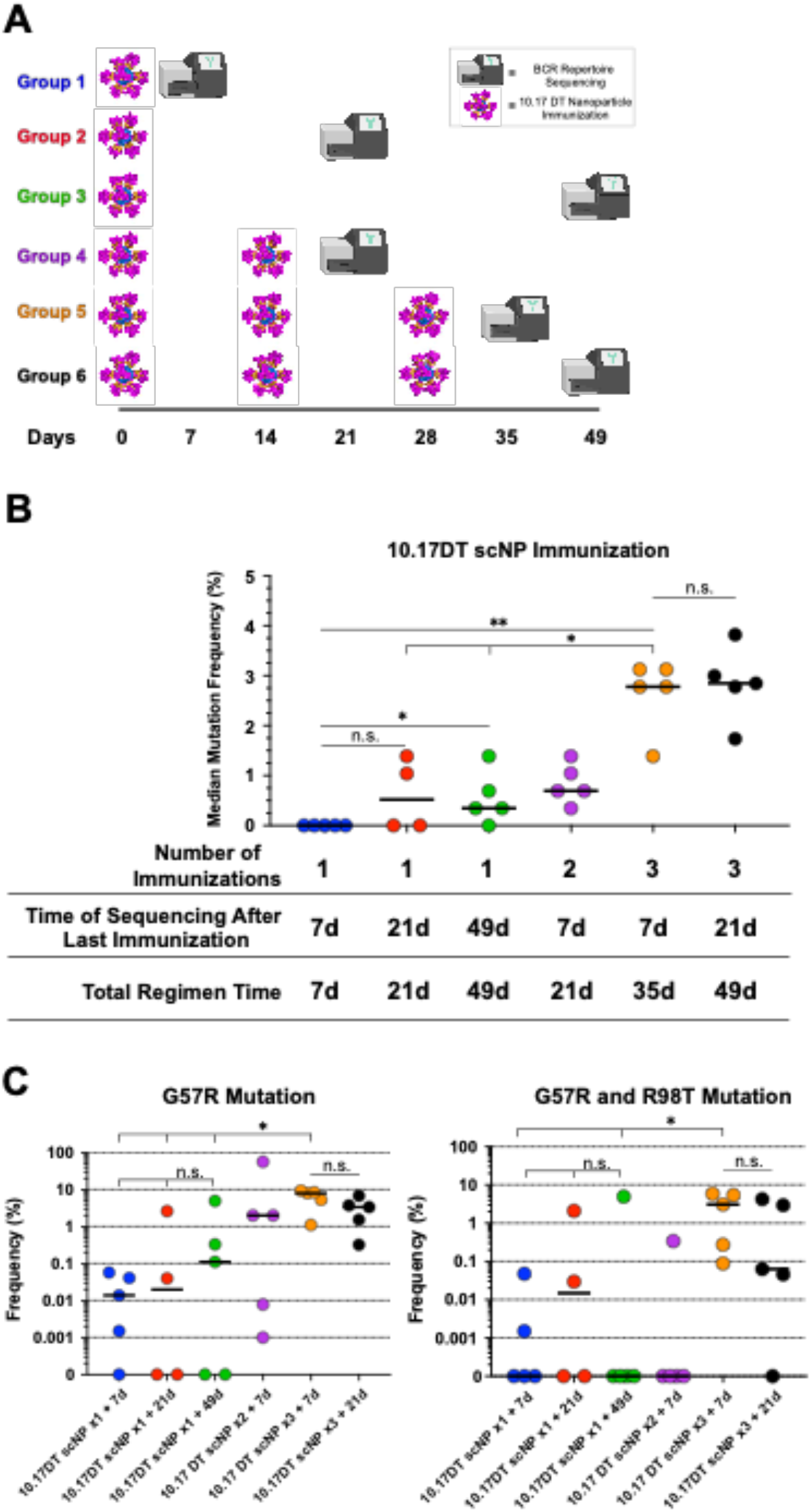
Priming with Multiple Immunizations Results in Higher Mutation Levels Than Single Immunizations with Prolonged Maturation Times. **A)** Immunization regimen and sequencing schedules of 10.17DT nanoparticle immunized mice (n=6 mice per group). **B)** Median somatic hypermutation frequency (computed as nucleotide mutations in V_H_ gene segment) of immunized mouse IgG heavy chain repertoires from NGS repertoire sequencing. C) Frequency of key early improbable mutations of DH270 UCA knock-in derived reads of immunized mouse IgG heavy chain repertoires or mouse light chain repertoires from NGS repertoire sequencing. Each dot represents the mutation frequency for the repertoire of one immunized mouse. * p-value <0.05, ** p-value <0.01, Wilcoxon-Mann-Whitney exact test. For visual clarity, bars denoting statistical significance testing are only shown for group comparisons referenced in the main text. See Data S3 for p values for all pairwise group comparisons. Bars denote group medians.

### Prime-boost regimen selects for combinations of key mutations

While frequent repetitive immunizations with a homologous priming immunogen initiates bnAb lineages in UCA knock-in mice (*6*), our data demonstrate homologous priming can result in diminishing returns on maturation progress beginning at about the third immunization. Once initiated with the V1-glycan deleted 10.17DT Env, the V3-glycan lineages like DH270 must eventually acquire accommodation of additional glycans on the V1 loop for development of neutralization breadth (*5*, *33*, *34*). We reasoned that presentation of a fully glycosylated V1 loop early in vaccination regimens would be critical to select against V1 loop-only directed lineages and select for combinations of functional improbable mutations. Thus, we designed a boosting immunogen with the V1 N133 and N138 glycan sites restored which we refer to here as 10.17 wildtype (WT).

We and others have proposed that utilizing immunogens with affinity differences to bnAb clone BCRs is important for guiding bnAb B cell lineage affinity maturation(*3*, *6*, *35*). We observed an affinity of 10.17WT that was over one order of magnitude higher for the DH270 UCA expressing both the G57R and R98T mutations than the DH270 UCA expressing the G57R mutation alone. We hypothesized that this affinity difference could provide a selective advantage for GCB cells bearing the G57R and R98T in combination (**Figure S4A**). We therefore chose 10.17WT with a fully glycosylated V1 loop that must be accommodated by mature V3-glycan bnAbs as a boosting immunogen.

We immunized mice 3 times with the 10.17DT nanoparticle and then boosted mice 5 times with 10.17 WT produced as a stabilized Env trimer (**Figure 4A**). We observed higher levels of heavy chain SHM in the prime-boost group compared to mice that only received the priming immunogen (p<0.0001; Wilcoxon-Mann-Whitney exact test) (**Figure 4B**). BCR repertoire sequencing showed a trend towards increased frequencies of individual key mutations G57R, R98T, L48Y as well as an order of magnitude higher frequency of the G57R and R98T combination in mice that received the prime-boost regimen compared to mice that received just the priming regimen alone (median mutation frequencies of the G57R and R98T combination of 0.351 and 3.61 for prime-only and prime-boost groups, respectively) (**Figure 4C, Figure S4B**). The key improbable light chain mutation S27Y was observed at a very low frequency in the BCR repertoires of both groups indicating limited selection of this critical early-stage maturation mutation by both the prime and prime-boost regimens using protein immunogens (**Figure S4B**). While serum neutralization of heterologous isolate 92RW020 trended higher in the prime-boost group (**Figure S4C**), the low titers of heterologous neutralizing antibodies in the serum of both immunization groups suggested maturing DH270 GC B cells derived with key improbable mutations had limited differentiation into plasma cells.

**Figure 4.**
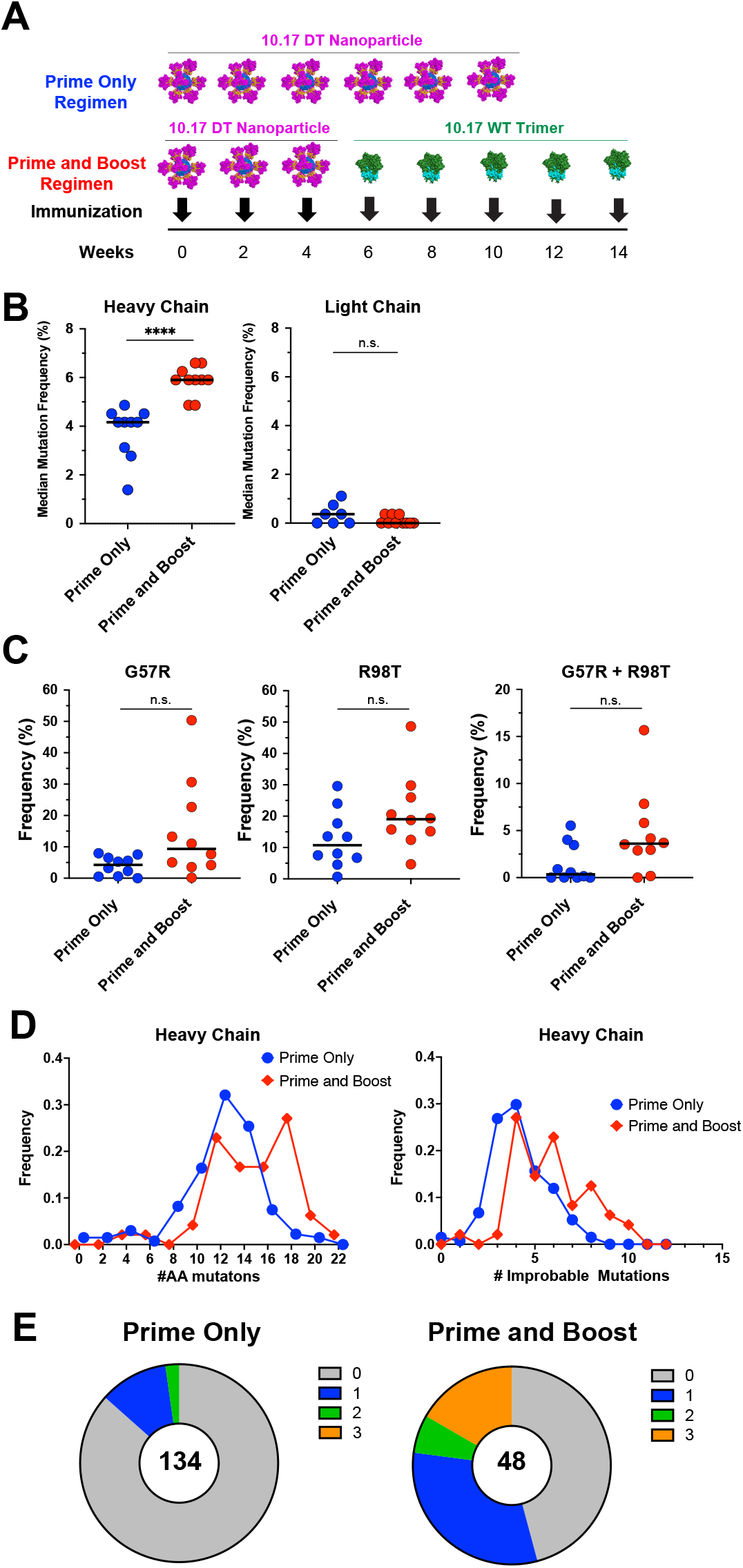
Prime-boost regimen selects for combinations of key improbable mutations. **A)** Immunization regimens for 10.17DT nanoparticle prime only (n=10 mice) and 10.17DT nanoparticle prime, 10.17 WT trimer boost mouse groups (n=12 mice). **B)** Median mutation frequency (computed as nucleotide mutations in V_H_ gene segment) of DH270 heavy and light chain reads (left two panels) and mean number of heavy and light chain improbable amino acid mutations (right two panels). **C)** Percentage of DH270 heavy chain reads in immunized mouse repertoires with key improbable mutations G57R (left), R98T (middle) or G57R and R98T in combination (right). **D)** Histograms of number of heavy chain amino acid mutations (left) and number of improbable amino acid mutations (right) in monoclonal antibodies isolated by antigen-specific single B cell sorting from the prime only (blue curves) and prime-boost (red curves) mouse groups. Each dot represents one immunized mouse in panels C and D. **E)** Portion of total vaccine-induced monoclonal antibodies that possess 0-3 of the four key improbable mutations (G57R, R98T, S27Y and L48Y) from mice receiving the prime only (left) or prime and boost (right) regimens. NGS repertoire sequence data from 5 mice in the prime-only group in panels B-D and monoclonal antibody data shown for the prime only group in panel E were originally reported in Saunders et al. (*6*). **** p-value <0.0001, ** p-value <0.01, Wilcoxon-Mann-Whitney exact test. Bars denote group medians.

Next, we performed paired heavy and light chain sequencing of 10.17DT-specific splenic B cells from the prime-boosted mice, and determined the frequency of combinations of critical mutations that co-occurred on both DH270 clonal lineage heavy and light chains. We observed significantly higher numbers of heavy chain mutations in antibodies isolated from the prime-boost immunized mice than in antibodies isolated from the prime-only immunized mice (p<0.0001; Wilcoxon-Mann-Whitney exact test) (*6*) (**Figure 4D**). Antibodies from the prime-boosted mice also acquired significantly higher numbers of improbable mutations than the prime-only mice (p<.0001; Wilcoxon-Mann-Whitney exact test) (**Figure 4D**). Because improbable mutations do not occur frequently by intrinsic AID activity prior to antigenic selection, these data demonstrated that the heterologous prime-boost regimen exerted stronger selective pressure on the DH270 B cell lineage than homologous priming alone. In addition, we observed higher numbers of specific key mutations in antibodies from the prime-boosted mice. A majority (54%; 26 out of 48) of the antibodies from the prime-boost group had at least one of the four key improbable mutations (G57R, R98T, S27Y, and L48Y) that we found were critical to the early acquisition of heterologous neutralization breadth in the DH270 lineage. This contrasted with what we previously observed in prime-only mice where most isolated monoclonal antibodies had not acquired any of the four improbable mutations (*6*) (**Figure 4E**). As an indication of how much further antibodies from the prime-boost group had matured, 17% of the antibodies had acquired 3 out of 4 key mutations. Thus, the heterologous prime-boosting regimen of 10.17DT followed by 10.17WT can select for combinations of the identical critical mutations *in the same B cell* as was also observed in the early development of the DH270 lineage in the setting of HIV-1 infection (*5*). Only a limited number of monoclonal antibodies acquired the S27Y mutation (**Figure S5**) indicating that selection of this mutation remains a substantial hurdle for protein vaccine regimens.

To determine if the success of the regimen was dependent upon the specific priming and boosting immunogens used, or if merely a product of 8 immunizations, we performed a second immunization study in which mice received the same 3x primes of the 10.17DT nanoparticle but were then administered 5 boosting immunizations with a different Env trimer for each boost. The 5 boosting immunogens were selected based on autologous virus neutralization sensitivity to DH270 antibody intermediates (*5*). Heavy chain somatic hypermutation was significantly lower (p <0.001; Wilcoxon-Mann-Whitney exact test) in the mice that were boosted with a series of different Env trimers than mice that were repetitively boosted with 10.17WT Env (**Figure S6**), demonstrating that boosting specifically with 10.17WT Env was critical to the success of the prime-boost regimen. Moreover, that an Env regimen not explicitly designed to target selection to key improbable mutations resulted in off-track maturation responses suggests that adoption of a mutation-guided vaccine strategy can be a more successful approach.

### Prime-boost regimen elicits bnAbs indicative of lineage maturation progress

Having elicited antibodies with combinations of key improbable mutations with a prime-boost regimen, we next sought to compare their binding and neutralization profiles to DH270.6 lineage members to monitor functional maturation progress. We produced a selection of recombinant antibodies from the mouse vaccine studies that acquired either 2 or 3 of the 4 key mutations (**Figure 5A**) and tested their binding against a set of 7 heterologous Env SOSIP trimers (**Figure S7A**). All vaccine-induced antibodies bound to the Envs bound by the first intermediate in the DH270.6 lineage (I5.6) and most vaccine-induced antibodies bound to 4 out of the 5 Envs bound by the second intermediate (I3.6). Recognition of heterologous Envs was therefore consistent with maturation progress past the first DH270 intermediate antibody and nearly reaching the second intermediate in the lineage. We next profiled the neutralization of the vaccine-induced antibodies using a panel of 24 viruses shown to be sensitive to DH270.6 neutralization (**Figure 5B**). In this HIV-1 isolate panel, the first intermediate (I5.6) in the DH270.6 lineage neutralized 12 out of 24 viruses. All of the vaccine-induced antibodies with 3 out of the 4 key mutations neutralized these same 12 viruses and did so more potently than the first intermediate. The second intermediate (I3.6) neutralized 18 of 24 viruses. Three of the vaccine-induced antibodies (DH270.MU89, DH270.MU90, and DH270.MU94) neutralized 17 out of the 18 viruses neutralized by the second intermediate I3.6. Thus, the neutralization breadth and potency of the vaccine-induced antibodies surpassed that of I5.6 and was approaching that of the I3.6 antibody (**Figure 5B**). We conclude that selection by our prime-boost regimen of 3 out of 4 critical improbable mutations resulted for the first time in the reproducible elicitation of bnAbs in a V3 glycan bnAb UCA knock-in mouse model with an unmutated CDRH3.

**Figure 5.**
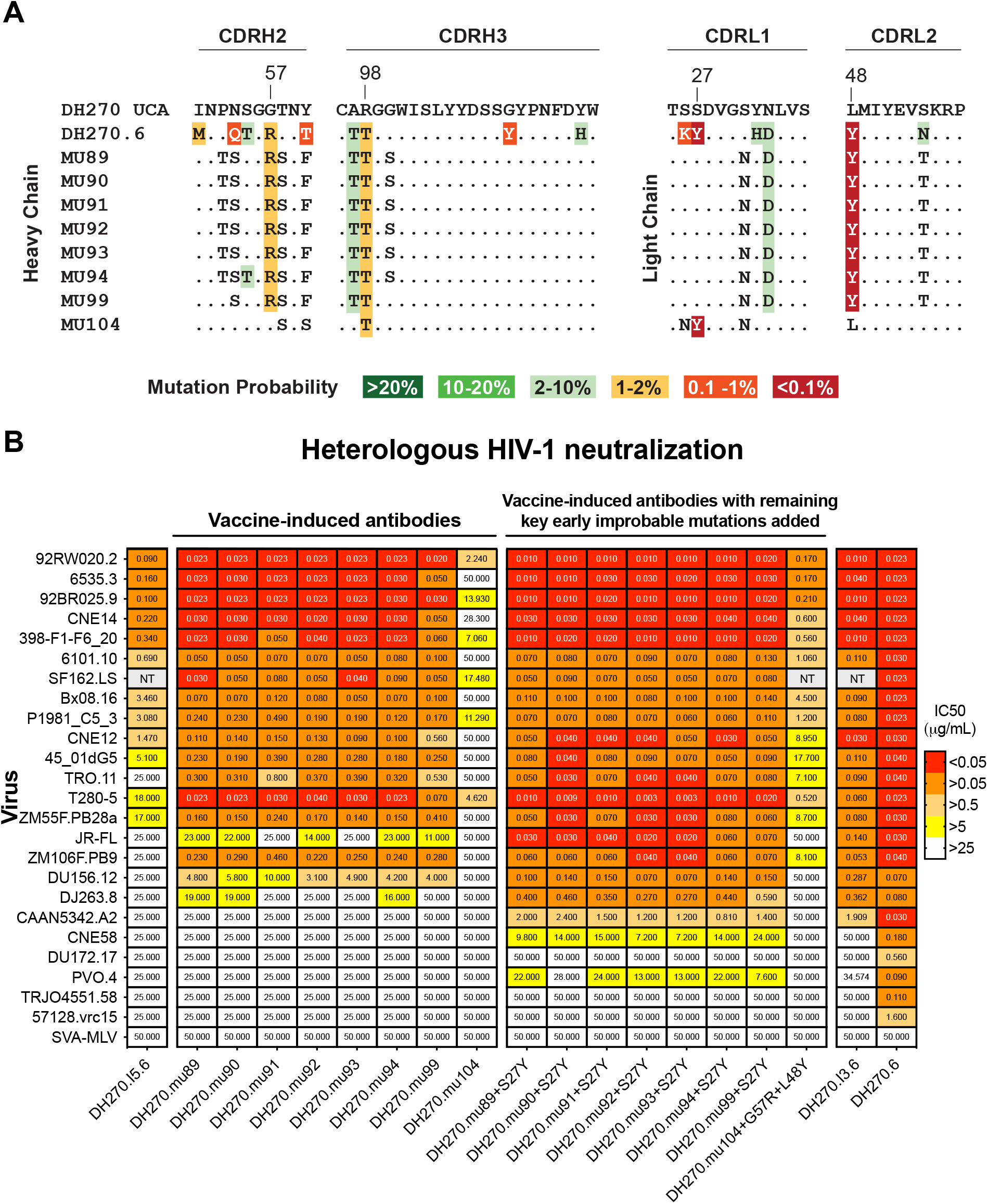
Binding and neutralization of prime-boost vaccine-induced monoclonal antibodies. **A**) Neutralization titers (ID50 values; mcg/ml) of prime-boost vaccine-induced antibodies against a panel of DH270.6 sensitive viruses. **B**) Alignment of prime-boost vaccine-induced antibodies to DH270 UCA and DH270.6 showing the acquisition of shared mutations with DH270.6. Mutations in vaccine-induced antibodies that are shared with DH270.6 are highlighted by their mutation probability in the absence of antigenic selection as estimated by the ARMADiLLO program. Dots in alignment represent amino acids at each position that match the reference sequence, DH270.UCA. Only select regions of interest (CDRH2, CDRH3, CDRL1, CDRL2) are shown. See Figure S5 for full length amino acid sequence alignments.

We next asked whether the addition of S27Y to vaccine-induced antibodies would result in further gains in neutralization breadth. We added the S27Y mutation to 7 vaccine-induced antibodies and observed improved neutralization breadth and potency in all mutant antibodies (Figure 5B). Antibody DH270.MU89+S27Y potently neutralized all 6 of the additional viruses neutralized by I3.6 and demonstrated increased breadth over I3.6 by neutralizing two additional viruses that were resistant to both I3.6 and DH270.key4 neutralization (**Figure 5B and 1C**). The observation of gains in neutralization breadth beyond what was observed in DH270.key4 suggested that mutations additional to the G57R, R98T, S27Y and L48Y mutations that were acquired by the vaccine-induced antibody were beneficial. We observed an improved affinity of DH270.MU89+S27Y relative to DH270.MU89 for heterologous Envs binding, but no improvement in binding affinity of DH270.MU89+S27Y to the 10.17DT or 10.17WT Envs (**Figure S7B**). The lack of a strong affinity gradient of 10.17DT and 10.17WT for S27Y was consistent with the limited selection of S27Y we observed in mice immunized with these protein immunogens. Together, these data indicated that S27Y is a key mutation for early-stage development of DH270-like antibodies and thus remains a high-value target for selection with additional boosting strategies.

### Structural Basis for Broad Neutralization by Prime-Boost Vaccination

To understand the structural basis for broad HIV-1 neutralization by a prime-boost regimen-induced antibody that had acquired critical improbable mutations and to define the structural requirements for boosting immunogens, we used single particle cryo-electron microscopy (cryo-EM) to determine the structure of the MU89 antigen-binding fragment (Fab) bound to the 10.17DT SOSIP trimer to an overall resolution of 3.6Å (**Figures 6, S8, and S9; Table S1**). The MU89 Fab bound to the Env V3-glycan site and utilized glycans N332 and N301, as well as the V1 and V3 loops for its paratope (**Figures 6 and S9**). The orientation of the Env-bound MU89 resembled that of the mature DH270.6 antibody, with the MU89 HCDR3 region similarly engaging the base of the gp120 V3-loop and glycan N332 (**Figure 6B, S9D and S9F**).

**Figure 6.**
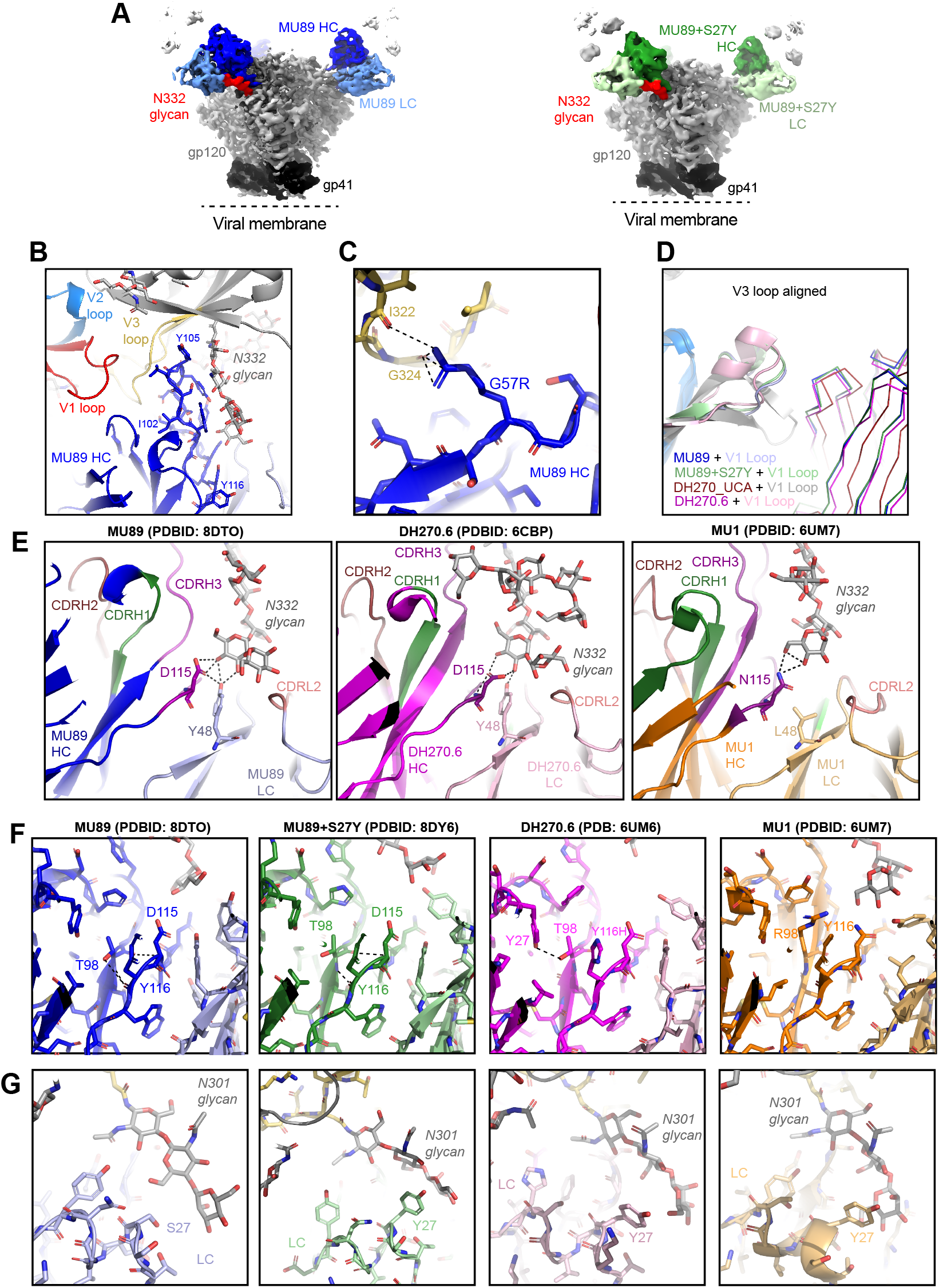
Structural mechanism of Broad Neutralization by a heterologous-prime boost vaccine induced antibody MU89. **A**) Left, cryo-EM reconstruction (3.6 Å resolution) of MU89 bound to 10.17DT SOSIP Env trimer segmented by components. Right, same as (left) but for the MU89+S27Y complex (4.3 Å resolution).**B**) Zoomed in view of MU89 Heavy chain (blue), binding interface with gp120. **C**) MU89 G57R binding conformation with V1 loop and GDIK motif. **D**) Overlay of MU89 (blue), MU89+S27Y (green), DH270.UCA (maroon), and DH270.6 (magenta) shown in lines with their corresponding V1 loops shown in cartoon. Structures were aligned by the V3 loop. **E**) Zoomed in views of heavy chain residue 115 and light chain residue 48 interacting with the N332 glycan. Polar interactions are shown in black dashed lines. **F**) Zoomed in views of key improbable mutation R98T interactions with surrounding residues. **G**) Antibody interactions with S27Y to glycan N301.

We had previously showed that antibody DH270.mu1 (referred to as MU1), induced in a prime-only regimen in a DH270 UCA knock-in mouse, bound Env with the gp120 V1 loop adopting an orientation conducive to the accommodation of V1 loop diversity (*6*). This V1 loop conformation allowed access of V_H_ residue R57, introduced in the antibody HCDR2 region due to the G57R improbable mutation, to the conserved V3 loop GDIR/K motif. The conformation of the MU89-bound V1 loop was similar with residue R57 engaging Env residues I322 and GDIK motif residue G324 (**Figure 6C and 6D**).

To understand the differences between Env recognition by MU1 and MU89 that led to the superior neutralization capability of MU89, we focused our attention on the key improbable mutations, V_L_ L48Y and V_H_ R98T, that occurred in MU89 but not in MU1. The L48Y and R98T improbable mutations together orchestrate local remodeling of the interactive surface for glycan N332 resulting in improved interactions with the glycan, as well as interactions that stabilize the internal structure of the antibody (**Figure 6E and 6F**). The side chain hydroxyl of V_L_ Y48 in DH270.6 and MU89 makes hydrogen bonding interaction with glycan N332, and with the side chain of V_H_ D115 (**Figure 6E**). By contrast, MU1 V_L_ L48 cannot similarly contact the N332 glycan, although the side chain of V_H_ N115 in MU1 hydrogen bonds with the terminal glycan N332 moiety. In DH270.6 and MU89, the V_H_ T98 side chain hydroxyl makes hydrogen bonding interactions with adjacent polar moieties including the backbone carbonyl and nitrogen of V_H_ H116 (Y116 in MU89), the backbone nitrogen of V_H_ D115, and a water mediated interaction with residue D115 side chain (**Figure SF and S9G**). While the resolution of the cryo-EM reconstructions precludes observation of water molecules, this water-mediated interaction is defined in the crystal structure of DH270.6 bound to the Man-V3 glycopeptide (PDB ID: 6CBP), (**Figure S9G**) and is likely to be conserved in the MU89 structure that harbors both the R98T substitution and the D115 residue that engage this water molecule.

We also determined a cryo-EM structure of antibody MU89+S27Y in complex with 10.17DT SOSIP trimer at an overall resolution of 4.32Å (**Figures 6A, S8, and S9**). The overall binding angle of MU89+S27Y closely resembled that of MU89 (**Figure S9F**). As was observed in the cryo-EM structures of the DH270.6-bound and MU1-bound complexes, the V_L_ S27Y substitution in MU89+S27Y resulted in the residue V_L_ Y27 side chain stacking against glycan N301, contributing additional binding contacts and likely stabilizing the glycan N301 conformation, thereby facilitating antibody access and binding to the V3-glycan site (**Figure 6G and S9C**).

Thus, critical interactions including essential light chain glycan contacts formed early in B cell maturation can confer broad recognition of the V3 glycan site. Moreover, vaccine regimens that can initiate rare V3 glycan bnAb precursors and then select for a few essential mutations necessary to counteract the Env glycan shield can achieve broad neutralization without needing to elicit the extremely high mutation frequencies (15-40%) typically observed in bnAbs from HIV infection.

### Nucleoside-modified mRNA-LNP prime-boost regimen selected for improbable glycan-contacting mutations at higher frequency than proteins

Nucleoside-modified mRNA vaccine technology holds tremendous potential for addressing the many challenges that must be overcome to produce an effective HIV vaccine. We have previously demonstrated that designed nucleoside-modified mRNA can encode well-folded SOSIP Env trimers and well-folded ferritin-Env nanoparticles (11). We next investigated whether the use of lipid nanoparticle (LNP)-encapsulated nucleoside-modified mRNA (mRNA-LNP) immunization could enhance vaccine-induced DH270.6 V3-glycan bnAb B cell lineage development. To detect differences between protein and mRNA-LNP regimens, we pooled the results from DH270 UCA V_H_^(+,-)^/V_L_^(+,-)^ knock-in mice that received similar protein prime-only regimens (n=10 mice), similar mRNA-LNP prime-only regimens (n=12 mice), similar protein prime-boost regimens (n=18 mice), and similar mRNA-based prime-boost regimens (n=12 mice). For all pooled prime-only groups, mice received 6 primes of 10.17DT nanoparticles. For all pooled prime-boost groups, mice received 3-4 primes with 10.17DT nanoparticles and 3-5 boosts with 10.17WT trimers (see **Data S1** for details on the vaccine regimens of the pooled groups). We observed no significant differences in the frequencies of total heavy chain mutations, total light chain mutations, total improbable heavy chain mutations, or key early-lineage heavy chain mutations (G57R, R98T or their combination) between mRNA and protein in either the prime-only or prime-boost groups (**Figures 7A, S10A and B**). Additionally, we saw no significant differences between the mRNA-LNP and protein immunized mice for serum neutralization of the heterologous 92RW020 virus (**Figure S10C**). Thus, mRNA-based regimens elicited similar patterns of heavy chain mutations and similar neutralization activity as protein immunizations. However, we did observe a critical difference in the response to mRNA-LNP was in significantly higher frequencies of total light chain improbable mutations elicited in the mRNA prime-boost group versus the protein prime-boost group (p<0.001; Wilcoxon-Mann-Whitney exact test) (**Figure 7A**). Specifically, improbable light chain mutations S27Y and L48Y were each selected at significantly higher frequencies by the mRNA prime-boost regimens compared to the protein prime-boost regimens (p<0.001; Wilcoxon-Mann-Whitney exact test) (**Figure 7B, C**). The frequency of the S27Y mutation increased by over 30-fold (**Figure 7B**) suggesting that the previously identified developmental bottleneck of acquiring S27Y may be at least partially alleviated through stronger selection of this specific mutation by the mRNA prime-boost regimen. Thus, improved acquisition of key light chain improbable mutations in the mRNA immunized mice indicates a vaccine modality to acquire the fourth critical improbable mutation (S27Y) lacking from our protein vaccine-elicited antibodies.

**Figure 7.**
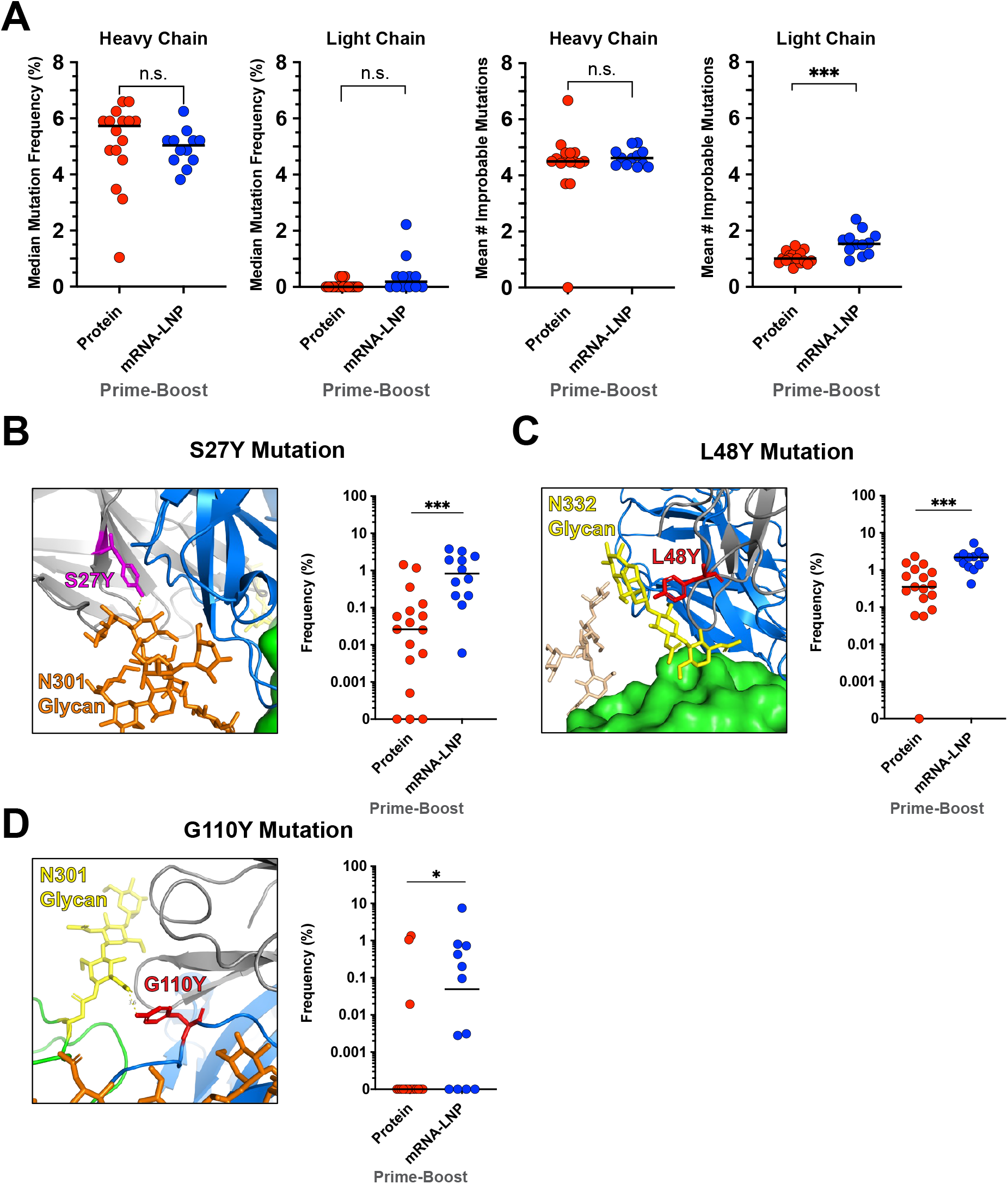
mRNA prime-boost immunization selects for higher numbers of V3 glycan-contacting improbable mutations. Groups of mice receiving similar mRNA-LNP or protein immunization regimens were pooled to compare elicited mutation frequencies and neutralization capacities. Details on regimens for the pooled mouse groups are listed in **Data S2**. **A)** Median mutation frequency (computed as nucleotide mutations in V_H_ or V_L_ gene segment) of DH270 heavy and light chain reads (left two panels) and mean number of heavy and light chain improbable amino acid mutations (right two panels) for pooled results from mRNA or protein primed and boosted mouse groups. mRNA immunization resulted in higher frequencies of V3 glycan contacting improbable mutations **B**) V_L_ S27Y, **C**) V_L_ L48Y and **D**) V_H_ G110Y. Frequency in the repertoire of DH270 reads containing glycan-contacting improbable mutations (B-D right panels) is shown beside structural views (B-D left panels; PDB: 6CBP) of the mutation’s interaction with V3 glycans. Each dot represents the repertoire of a single immunized mouse. * p-value <0.05, *** p-value <0.001, Wilcoxon-Mann-Whitney exact test. For visual clarity, bars denoting statistical significance testing are only shown for group comparisons referenced in the main text. See Data S3 for p values for all pairwise group comparisons. Bars denote group medians.

From the pooled groups, we next selected a pair of mouse groups administered similar prime-boost regimens to perform a side-by-side comparison of single mRNA vs. protein immunization experiments (**Figure S11A**). We observed similar levels of 10.17DT specific GC B cells (**Figure S11B**) and mutation patterns that were consistent with the those observed in the pooled groups including key light chain glycan-contacting improbable mutations selected at significantly higher frequencies in the mRNA group (**Figure S11C and D**). In this comparison of single groups of primed and boosted mice, we also observed higher repertoire diversity in the mRNA group (**Figure S11E**) and significantly higher neutralizing titers of vaccine-boost matched virus 10.17WT and of heterologous virus 92RW020 in the mRNA group (p<0.01; Wilcoxon-Mann-Whitney exact test) (**Figure S11F**) underscoring the potential of mRNA-based vaccines for eliciting superior antibody responses.

The increased frequencies of S27Y and L48Y observed in mRNA-LNP prime-boosting regimens led us to investigate why these two specific mutations were selected at higher frequencies. S27Y and L48Y are both highly improbable with an expected frequency of <0.1% prior to antigenic selection, with S27Y forming a pi-stacking interaction with the base of the N301 glycan and L48Y forming a hydrogen bond with a terminal mannose residue in the N332 glycan (**Figure 7B and C**) (*36*). Based on the role of S27Y and L48Y in glycan recognition, we next examined the selection of improbable heavy chain mutation G110Y which interacts at the base of the N301 glycan and is the only other glycan-contacting mutation in the set of 12 mutations of the DH270min minimally-mutated bnAb. Our analysis of pooled groups of protein and mRNA-LNP primed and boosted mice showed that, like S27Y and L48Y, the G110Y frequency was significantly higher in mRNA-LNP-immunized mice (**Figure 7D**) (p<0.05; Wilcoxon-Mann-Whitney exact test). The increased frequencies of the three most critical glycan-contacting mutations in the DH270 lineage is consistent with a superior capability of mRNA-based immunogens to select glycan-contacting mutations and suggests that HIV-1 vaccine strategies that optimize glycan presentation for the selection of hard-to-elicit glycan-contacting mutations may be critical to successful induction of bnAbs.

## DISCUSSION

In this study, we have demonstrated the success of a process for design of sequential immunizations that can select for the accumulation of key improbable mutations in bnAb B cell lineages. Acquisition of four early-lineage improbable mutations resulted in 79% of the neutralization capacity of a V3-glycan bnAb B cell lineage. We used these four mutations as markers to monitor early maturation progress in V3-glycan bnAb UCA knock-in mice, investigated the kinetics of their acquisition in repetitive immunization regimens, and showed a plateaued response in their acquisition after 3 priming immunizations. The frequency of mutations attained was not explained by prolonged maturation times but rather was likely indicative of continued antigenic fueling of the germinal center by repetitive immunization over short time intervals. We demonstrated the successful design of a prime-boost regimen for V3-glycan B cell lineage maturation and showed that this regimen, using either protein or nucleoside-modified mRNA-LNP immunogens, can select for higher frequencies of improbable mutations over those selected by the prime immunization alone. In addition, we isolated vaccine-induced V3-glycan bnAbs and demonstrated the structural basis of neutralization breadth induced by the prime-boost combination. Finally, we showed that nucleoside-modified mRNA-LNP prime-boost regimens were superior in selection of improbable glycan-contacting mutations compared to protein immunizations.

Since bnAbs develop in persons living with HIV-1 only after years of prolonged viremia and viral diversification (*5*, *14*, *37*), the goal of HIV-1 vaccine development is to design regimens to mimic continuous viral antigen exposure with envelope variants that can select for rare B cells that can lead to bnAb breadth (*3*). Repetitive immunizations, escalating doses of immunogen over multiple weeks and implanted pumps with continuous antigen administration can all increase Env immunogenicity and prolong antigen stimulation in B cell GCs (*38*–*41*). However, the most critical parameter of HIV-1 vaccination regimens is the choice of boosting Envs capable of selecting improbable mutations that confer broad neutralization in bnAb B cell lineage BCRs (*4*–*6*, *19*). We and others have designed or identified immunogens for stimulating and expanding bnAb precursors for CD4 binding site (*35*, *42*–*45*), V2-glycan (*8*, *46*–*48*), V3-glycan (*6*, *20*, *49*, *50*) and gp41 membrane proximal external region bnAb B cell lineages (*26*). For sequential HIV-1 vaccine regimens to reach the ultimate goal of bnAb induction, they must build off of the recent successes in designing priming immunogens by effectively guiding B cell maturation to acquire the critical improbable mutations necessary for broad heterologous virus neutralization. Central to this effort is the design of boosting immunogens to provide higher affinity binding to antibodies with desired mutations than antibodies without desired mutations which provides a selective advantage to B cells evolving toward bnAb activity (*3*, *6*, *35*, *51*). Also key is the use of immunogens that can select for bnAb B cell maturation events that allow lineage members to bind and/or accommodate Env glycans. Accommodation of the V1 loop is a critical milestone in the generation of V3-glycan-targeted bnAbs (*5*, *6*, *33*, *34*, *52*). Here we used an Env with V1 glycans deleted as a prime (*6*) and an Env with the V1 glycans restored as a boost to select for V3-glycan bnAb lineage improbable mutations. Coupled with the affinity gradient observed with the boosting Env for antibodies with functional improbable mutations, this prime and glycosylated Env-boost combination resulted in induction of antibodies with considerable heterologous breadth.

Strategies to accelerate bnAb induction in the vaccine setting are clearly needed given the process typically takes multiple years and exposure to potentially billions of antigenic variants with only limited success rates in the setting of natural infection. Here, through the adoption of a mutation-guided vaccine design strategy, we have shown that the precise mapping of improbable mutations in a bnAb lineage can be used to identify rate-limiting steps in bnAb maturation, that boosting immunogens can be designed to overcome those steps, and that B cell responses to vaccination with sequential prime-boost regimens can be evaluated via BCR repertoire sequencing by measuring the frequency by which those rate-limiting mutations are acquired. Much in the same way that an enzyme catalyzes a reaction, the specific design of boosting immunogens that target mutation selection to speed up rate-limiting steps in bnAb development may be an effective strategy to accelerate bnAb induction. While improbable mutations can arise after a single immunization (*53*), it is only after the selection of combinations of the key improbable mutations that confer neutralization breath can bnAb B cell induction occur. Thus, the essential targets for selection by an HIV-1 vaccine should be combinations of *functional* improbable mutations. We have shown here that defining the functional improbable mutations in a lineage, designing boosts that can select for these mutations, and monitoring their acquisition in immunized animal models can be an effective strategy for developing an HIV-1 vaccine.

It is also important to point out that design strategies aimed solely at generating boosting immunogens with affinity gradients to intermediate antibodies could lack the precision required to directly select for functional improbable mutations. The design of immunogens for binding with high affinity to intermediate antibodies could preferentially select for the non-essential mutations at the expense of the functional mutations, thereby omitting critical somatic mutations required for on-track maturation toward bnAb activity. The mutation-guided vaccine design strategy increases precision by making selection of functional improbable mutations a specific design criterion for boosting immunogens, thus providing on-track B cells with a selective advantage.

We demonstrated proof-of-principle that the mutation-guided design strategy can direct maturation towards bnAb activity in a mouse model that expresses a pre-rearranged UCA, yet work is still needed to adapt this strategy to a setting with both more diverse and more limited frequency precursors. Previous studies have demonstrated convergent maturational events among bnAbs within the same epitope class (*2*, *4*, *16*, *54*) which suggests that by adjusting the design parameters of the mutation-guided design strategy one could design boosting immunogens that select for these types of shared mutations more generally.

Nucleoside-modified mRNA-LNP is distributed to multiple immune organs after intramuscular (IM) immunizations(*55*), and express *in vivo* over 10-14 days, thus prolonging immunogen delivery and GC loading over IM protein administration (*56*). Protein prime-boost immunizations selected for three of the four key improbable mutations for DH270 V3-glycan bnAb breadth, but only rarely selected for key improbable light chain mutations required for bnAb binding to glycans at N301 or N332. Here, the administration of mRNA-LNP encoding prime-boost immunogens identical to those administered as proteins were superior for selection of key improbable mutations required for glycan binding. Based on the superior selection of the three most critical glycan contacting mutations in the DH270 lineage, we hypothesize that mRNA-expressed immunogens, which are mainly produced by host dendritic cells, may exhibit glycosylation patterns that are more favorable for selecting key improbable glycan-contacting mutations than recombinantly produced proteins. Glycan engineering of Env-based immunogens by the complete removal of glycan sites, the systemic modification of glycoforms by enzymatic treatment, or by production in glycosylation machinery-deficient cell lines, has been the most widely used strategy for designing HIV-1 priming immunogens to date (*6*, *20*, *21*, *35*, *42*–*45*, *50*, *57*). However, tailoring immunogens to express specific glycoforms at targeted sites to select critical glycan-contacting mutations is very difficult. Thus, we are encouraged that mRNA-based immunogens could provide more optimal glycosylation presentation than recombinantly produced protein immunogens. As the current mRNA-LNP vaccine regimens for SARS-CoV-2 demonstrate, mRNA-LNP can increase neutralizing Ab affinity and breadth (*58*). Similarly, our data indicate that use of mRNA-LNP prime and boosting regimens may be a mode of increasing the efficacy of HIV-1 Env prime boost vaccine regimens. An additional strategy may be to design a vaccine that uses both mRNA-LNPs and proteins as either primes or boost immunogens. Such a vaccine design may capitalize on the durability of protein immunization, while also taking advantage of the improved selection of key improbable bnAb light chain mutations by mRNA-LNP formulation.

V3-glycan bnAb precursors are rare (*20*) and their maturation pathways are punctuated with numerous improbable mutations (*4*), many of which are required for full bnAb activity (*20*, *27*). Thus, the remaining hurdles to induction of bnAbs include priming of sufficient numbers of rare V3 glycan bnAb precursors in humans, iterative design of subsequent Env boosts to select for the remainder of functional bnAb mutations required for full bnAb neutralization breadth, and the generation of robust and durable serum bnAb responses.

Finally, these studies demonstrate a generalized process of HIV-1 vaccine immunogen development whereby boosting immunogens are chosen based on their ability to select for key functional improbable mutations. Critical to further success will be to implement such a process of immunogen design in a high throughput mode to design Envs that can select for key mutations in fully matured bnAbs, and to do so in many bnAb B cell lineages against multiple Env bnAb epitopes (*2*).

## Acknowledgments

We thank Austin Harner, Courtney Peninger, and Justine Mae Flores for veterinary assistance; Bhavna Hora and the DHVI Viral Genetics Analysis Core facility for mouse repertoire sequencing; Brian Watts for technical assistance with binding kinetics assays; Elizabeth Donahue, Kelly Cuttle, and Whitney Edwards for program management. We thank Zekun Mu and RJ Edwards for helpful discussions. Flow cytometry and FACS were performed in the Duke Human Vaccine Institute Flow Cytometry Shared Resource, and we thank Letealia Oliver for technical assistance. Cryo-EM data were collected at the Shared Materials Instrumentation Facility at Duke University as part of the Molecular Microscopy Consortium.

## Author contributions

Conceptualization: KW, KOS, BFH

Investigation: KW, KOS, VS, DWC, SV, JSMB, MadB, TE, SMX, CJ, CB, XL, MEB, JB, AS, HC, AE, MAT, CBF, YT, CB, MatB, HM, SMA

Supervision: KW, AN, BH, DCM, FWA, DW, WBW, KOS, PA, BFH

Data Analysis: KW, KOS, PA, VS, DWC, SV, JSMB, MB, BFH

Funding acquisition: BFH

Writing – original draft: KW, KOS, BFH

Writing – review & editing: All authors

## Data and materials availability

The protein structures of DH270.MU89 and DH270.MU89+S27Y have been deposited in the PDB (XXXX and XXXX). Electron microscopy data has been deposited under accession numbers: XXX. Vaccine-induced antibody sequences DH270.MU89-94, DH270.MU99, and DH270.MU104 have been deposited in GenBank under accession numbers XXX. Mouse BCR repertoire heavy and light chain NGS datasets have been deposited at SRA under accession numbers XXX. All flow cytometry data and binding kinetics data are available upon request. All other data has been made available in the main text or supplementary materials. The DH270 UCA VH/VL KI mice and CH235 UCA VH/VL KI mice are available from F.W.A.’s laboratory under a standard material transfer agreement with Boston Children’s Hospital. 3M-052 stable emulsion adjuvant is available from M. Tomai under a material transfer agreement with 3M Company (St. Paul, MN).

## Supplementary Materials

### Materials and Methods

Fig. S1. Neutralization of antibody mutants with all combinations of 5 mutations from the second intermediate I3.6 introduced into the first intermediate I5.6 of the DH270.6 clonal lineage Fig. S2. Mutation frequencies and neutralization profiles in response to repetitive immunizations Fig. S3. Mutation acquisition dynamics in response to single immunizations with varying maturation times vs. repetitive immunization with the 10.17DT nanoparticle.

Fig. S4. Binding kinetics of prime-boost immunogens, light chain mutation frequencies, and neutralization of primed and boosted mice

Fig. S5. Prime-boost immunization elicited antibodies with somatic mutations shared with DH270.6

Fig. S6. Affinity Maturation of Monoclonal antibodies elicited by 10.17DT nanoparticle prime and 10.17WT SOSIP trimer boost regimen results in Broad Env Reactivity and high affinity for Heterologous HIV-1 envelopes.

Fig. S7. Affinity Maturation of Monoclonal antibodies elicited by 10.17DT nanoparticle prime and 10.17WT SOSIP trimer boost regimen results in Broad Env Reactivity and high affinity for Heterologous HIV-1 envelopes.

Fig. S8. Cryo-EM data processing for the MU89 and MU89+S27Y bound complexes to HIV-1 Env Trimer.

Fig. S9. Structural details of Env binding interfaces of vaccine elicited antibodies MU89 and MU89+S27Y.

Fig. S10. Highly similar mutational responses in mRNA-LNP and protein immunized mouse groups for both prime-only and prime-boost regimens.

Fig. S11. mRNA-LNP prime-boost regimen results in higher frequency of light chain improbable mutations and higher heterologous neutralization than protein prime-boost regimen.

Table S1. Cryo-EM data collection and refinement statistics

Data S1. mRNA and protein immunization studies used in analysis of pooled results

Data S2. Details of all mouse immunization regimens

Data S3. P-values of pairwise mouse group comparisons

