## Supplemental Material for "Mutation-Guided Vaccine Design: A Strategy for Developing Boosting Immunogens for HIV Broadly Neutralizing Antibody Induction"

**This PDF file includes:**

Materials and Methods
Supplementary Text
Figs. S1 to S11
Table S1

**Other Supplementary Materials for this manuscript include the following:**

Data S1 to S3

#### Materials and Methods

Mouse immunizations

DH270 UCA heterozygous heavy chain variable and light chain variable region double knock-in mice ($V_HD_HJ_H^{+/-}$, $V_\lambda J_\lambda^{+/-}$ KI strain) and CH235 UCA heterozygous heavy chain and light chain variable region double knock-in mice ($V_HD_HJ_H^{+/-}$, $V_\lambda J_\lambda^{+/-}$ KI strain) were generated as previously described (*6*). Briefly, DH270 UCA KI mice administered protein immunogens were immunized intramuscularly and/or subcutaneously (see **Data S2** for specific details of immunization regimens for each mouse group) with 25 mcg of protein immunogen along with 5 mcg of the TLR4 agonist-based adjuvant GLA-SE (AAHI). Repertoire sequencing data from a group of DH270 UCA KI mice primed with with 10.17DT scNP and previously described in Saunders et al. (*6*) were combined with an additional group of mice primed with 10.17DT scNP to increase the number of prime-only mice for group comparisons (see **Data S2** for specific details). Single antibody sequences from isolated monoclonal antibodies from the 10.17DT scNP prime-only mice from Saunders et al. were also used in comparisons to the antibody sequences from monoclonal antibodies isolated from the protein prime-boost regimen in Figure 4. CH235 UCA KI mice were immunized with 25 mcg of protein immunogens and 0.5 μg of 3M-052 aqueous formulation admixed with 50 μg of Alum in PBS (AAHI) at 2 sites (hind limbs) intramuscularly and 2 sites subcutaneously (fore limbs). Mice administered mRNA-LNP immunogens were immunized with 20 mcg of the mRNA-LNP immunogen. Mice were administered immunogens at 2 week intervals and necropsied 7 days after the last immunization unless otherwise noted. Full details of each immunization regimen including dose, adjuvant, route of immunization, immunization schedule, and time of specimen collection and sequencing for all groups of mice used in this study are listed in supplementary **Data S2**. All mice were cared for in a facility accredited by the Association for

Assessment and Accreditation of Laboratory Animal Care International (AAALAC). All study protocol and all veterinarian procedures were approved by the Duke University Institutional Animal Care and Use Committee (IACUC).

Nucleoside-modified mRNA-LNP production

Nucleoside-modified mRNA-LNP were produced as previously described (*59*). Briefly, sequences of furin and HIV Env constructs CH848.3.D0949.10.17chDS.SOSIP_N133DN138T-ferritin, CH848.10.17.N133DN138T E169K_DS_LSK ferritin 2x linker, CH848.3.D0949.10.17_D230N_H289N_P291S_E169K_DS.SOSIP and CH848.10.17.N133DN138T_D230N_H289N_P291S_E169K_DS_VRC ferritin were codon optimized, synthesized (GenScript), and cloned into mRNA production plasmids. After ligation into the expression vectors, mRNAs were produced using T7 RNA polymerase (MEGA-script, Ambion) on linearized plasmids. mRNAs were transcribed to contain 101-nt-long poly(A) tails. m1□□5'-triphosphate (TriLink) instead of UTP (uridine 5'-triphosphate) was used to generate modified nucleoside-containing mRNAs. The furin-encoding mRNA was capped using the ScriptCap m7G capping system and ScriptCap 2′-O-methyl-transferase kit (ScriptCap, CellScript) (*60*). Capping of *in vitro* transcribed HIV Env-encoding mRNAs was performed co-transcriptionally using the trinucleotide cap1 analog, CleanCap (TriLink). All mRNAs were purified by cellulose purification, as described (*61*). All mRNAs were analyzed by agarose gel electrophoresis and were stored frozen at -20 °C. Nucleoside-modified HIV-encoding mRNAs were encapsulated in LNP for mouse immunizations as previously described (*62, 63*). Modified mRNAs in aqueous phase were rapidly mixed with a solution of lipids dissolved in ethanol. LNP formulation contains ionizable cationic lipid (proprietary to Acuitas)/phosphatidylcholine/cholesterol/PEG-lipid. The cationic lipid and LNP composition are described in US patent US10,221,127 (*64*).

Nucleoside-modified mRNA transfection in 293F cell line

mRNA-encoded 10.17DT DS 2xGS sgNPs SOSIP-ferritin nanoparticles were expressed in 293F cells and purified as previously described (*59*). 293F cells were diluted to 0.7x$10^6$ cells/ml 24 h before transfection. On the next day, cells were diluted again to 1x$10^6$/ml and 120 ml of cells were seeded into 250 ml tissue culture flasks for transfection. 10.17DT 2xGS sgNP mRNAs were transfected at 0.4 μg/$10^6$ cells with the addition of 0.1 μg/$10^6$ cells of furin mRNAs. TransIT-mRNA Transfection Kit (Mirus Cat# MIR2250) was used for mRNA transfection following the manufacturer's instructions. Transfected cells were cultured at 37°C with 8% $CO_2$ and shaking at 120 rpm for 48 h (for gp160) or 72 h. 72 h later, 10.17DT sgNPs were purified from supernatant by *Galanthus nivalis* lectin (GNL) beads followed by size-exclusion chromatography.

Recombinant SOSIP envelope production

Soluble Env trimers were expressed in Freestyle 293F cells by transient transfection and purified by PGT145 or PGT151 affinity chromatography as previously described (*6*). Trimeric envelope was purified by size exclusion chromatography with a Superose6 16/600 column (GE Healthcare) in 10 mM Tris pH8, 500 mM NaCl. To produce biotinylated CH0848 10.17DT SOSIP gp140s, the envelope sequence was expressed with a C-terminal avidin tag (AviTag: GLNDIFEAQKIEWHE). 25 μM of envelope was biotinylated with the BirA biotin-protein ligase standard reaction kit (Avidity) for 5 h at 30°C. The biotinylated protein was then purified with a Superose6 16/600 column (GE Healthcare) in 10 mM Tris pH8, 500 mM NaCl. Fractions containing trimeric HIV-1 Env protein were pooled together, sterile-filtered, snap frozen, and stored at -80 °C.   Additionally, we produced an enhanced version of the 10.17DT envelope SOSIP trimer referred to as CH848 10.17DTe where autologous virus-specific immunogenic regions ("glycan holes") were occluded by adding glycans at sites N230 and N289 (by introducing mutations of D230N, H289N, P291S to 10.17 DT) in addition to adding the E169K mutation for restoring the V2 apex bnAb epitope.

Nanoparticle production

*Protein nanoparticles.* Ferritin nanoparticles presenting Env SOSIP trimers were produced and characterized as previously described (*6*).  CH848 10.17DT SOSIP gp140 was expressed with amino acids LPSTGG encoded at its c-terminus. The CH848 10.17DT SOSIP was expressed in Freestyle293 cells and purified by PGT151 affinity chromatography. Trimeric gp140 was isolated by size exclusion chromatography using a Superose6 16_60 column (Cytvia). Ferritin nanoparticles were expressed with a pentaglycine repeat sequence encoded at the n-terminus of each subunit. His-tagged ferritin nanoparticles were purified by HisTrap prepacked 5 mL column (Cytvia). Purification tags were HRV3C digested off of the ferritin molecule. CH848 SOSIP with a C-terminal sortase A tag and ferritin particles with a sortase A N-terminal tag were buffer exchanged into 50mM Tris, 150mM NaCI, 5mM CaCI2, pH7.5. 120 μM of SOSIP gp140 was mixed with 120 μM of ferritin subunits and incubated with 100 μM of sortase A overnight at room temperature. Envelope-conjugated nanoparticles were isolated by size exclusion chromatography using a Superose6 16_60 column in 10 mM Tris. 500 mM NaCl, pH8. Proteins were immediately snap frozen and stored at -80° C.

*mRNA-LNP nanoparticles.* For generating mRNA expressed nanoparticles presenting Env trimers, single gene constructs encoding ferritin subunits fused to gp140 SOSIP subunits were produced as previously described (*59*). Briefly, CH848 10.17DT SOSIP trimer-ferritin NPs were produced by gene fusion of CH848 10.17DT SOSIP trimer gene with the *Helicobacter pylori* (*H. pylori)* ferritin gene (*FtnA*) (GenBank NP_223316). We utilized three different versions of mRNA-LNP 10.17DT ferritin nanoparticles in separate immunization regimens. The first immunogen consisted of 10.17DT SOSIP fused to the fifth amino acid of wildtype *H. pylori* ferritin using a GGGSG linker. The second construct fused the gene of 10.17DTe to the second amino acid (Leu) of wildtype *H. pylori* ferritin using a GGGSGGGGSG (termed "2xGS") linker. The third construct fused the gene of 10.17DTe to the fifth amino acid of *H. pylori* ferritin with a glycine and a serine added to the C-terminus and an N19Q mutation, which removes an N-linked glycosylation site at position 19 (termed "VRC ferritin") (*65*).

Antigen-specific single B cell sorting and antibody isolation

Antigen-specific single B cell sorting, antibody isolation and antibody sequencing was performed as previously described (*6*). Memory B cells from splenocytes were stained and sorted by fluorescence activated cell sorting using a panel of fluorochrome-antibody conjugates (all from BD Biosciences): BB700 anti-mouse IgG1 (A85-1), BB700 anti-mouse IgG2a/2b (R2-40), BB700 anti-mouse IgG3 (R40-82), PE anti-mouse GL7 (GL7), PE-Cy7 anti-mouse IgM (R6-60.2), AlexaFluor700 anti-mouse CD19 (1D3), BV510 anti-mouse IgD (11-26C.2a), and BV650 anti-mouse B220 (RA3-6B2). Cells were also labeled with biotinylated SOSIP envelope trimers (10.17 or 10.17DT) conjugated to streptavidin-BV421 or streptavidin-AF647. Env-specific memory B cells were identified as viable B220+CD19+IgM-IgD-GL7-IgG1/2/3+ cells that bound both BV421- and AF647-conjugated SOSIP trimers. Single cells were sorted on a BD

FACS AriaII into 96-well PCR plates containing lysis buffer. Plates were immediately snap frozen and stored at -80° C.

Immunoglobulin genes were amplified as previously described with some modifications (*26, 66*). Immunoglobulin genes from a single B cell were reverse transcribed with Superscript III (ThermoFisher) using random hexamer oligonucleotides as primers. The complementary DNA was used to perform nested PCR for DH270 heavy and light chain genes using AmpliTaq gold (ThermoFisher) and primers designed to bind to the DH270 UCA variable region sequence. In parallel, PCR reactions were done with mouse immunoglobulin-specific primers. PCR amplicons were identified by gel electrophoresis and purified for Sanger sequencing using a PCR clean-up kit (Qiagen). Contigs of the PCR amplicon forward and reverse sequences were made, and immunogenetics annotation was performed with Cloanalyst(*67*) using both the human and mouse Ig gene libraries. Antibody genes were categorized as human or mouse. A second aliquot of the purified PCR amplicon was used for overlapping PCR to generate a linear expression cassette. The expression cassette was transfected with ExpiFectamine (ThermoFisher) into Expi293F cells (ThermoFisher). Clarified cell culture supernatant was analyzed by ELISA for binding to envelope. Antibodies with desired binding profiles were selected for gene synthesis and cloning of expression plasmids (GenScript).

B cell immunophenotyping

Immunophenotyping of murine B cells was performed as described previously (*6, 59*). Briefly, spleens from immunized mice one week after the last immunization (unless otherwise noted; see **Data S2** for individual mouse study details including specimen collection times) were processed into single-cell suspensions and treated with ACK lysis buffer to remove red blood cells.

Splenocytes were incubated with optimized concentrations of fluorochrome-mAb conjugates FITC anti-mouse IgG1 (A85-1), FITC anti-mouse IgG2a/2b (R2-40), FITC anti-mouse IgG3 (R40-82), PerCP-Cy5.5 CD21 (7E9), PE anti-mouse GL7 (GL7), PE-CF594 anti-mouse CD93 (AA4.1), PE-Cy5 anti-mouse CD38 (90), PE-Cy7 anti-mouse IgM (R6-60.2), AlexaFluor700 anti-mouse CD19 (1D3), BV510 anti-mouse IgD (11-26C.2a), BV570 anti-mouse CD11b (M1/70), BV605 anti-mouse CD95 (Jo2), BV650 anti-mouse B220 (RA3-6B2), BV711 anti-mouse CD138 (281-2), and BV786 anti-mouse CD23 (B3B4). Dead cells were identified by labeling with Near-IR Live/Dead (Thermo Fisher Scientific).Cells were analyzed on a BD LSRII (BD Biosciences). Data were analyzed using FlowJo v10 (FlowJo).

In vitro HIV-1 neutralization

Antibody-mediated HIV-1 neutralization was measured using Tat-regulated luciferase (Luc) reporter gene expression to quantify reductions in virus replication in TZM-bl cells as described previously(*68*). TZM-bl cells were obtained from the NIH AIDS Research and Reference Reagent Program, as contributed by John Kappes and Xiaoyun Wu. The monoclonal antibody or serum was pre-incubated with virus (~150,000 relative light unit equivalents) for 1 h at 37 °C, and TZM-bl cells were subsequently added. After 48 h cells were lysed and Luc activity measured using a GloMax Navigator luminometer and Brite-Glo Reagent (Promega). Neutralization titers are the inhibitory concentration at which relative luminescence units (RLU) were reduced by 50% or 80% compared to RLU in virus control wells after subtraction of background RLU in cell control wells (IC50 and IC80 respectively).

Cryo-EM data collection, processing, and model fitting

To prepare Env complexes, CH848 10.17DT SOSIP trimer at a final concentration of ~2 mg/mL was incubated with 4- to 6- fold molar excess of the MU89 or MU89+S27Y Fabs for 30 to 60 min. To prevent aggregation during vitrification, the sample was incubated in 0.085 mM dodecyl-maltoside (DDM). A 2.5-µL drop of protein was deposited on a Quantifoil-1.2/1.3 grid (Electron Microscopy Sciences, PA) that had been glow discharged for 10 seconds using a PELCO easiGlow™ Glow Discharge Cleaning System. After a 30-second incubation in >95% humidity, excess protein was blotted away for 2.5 seconds before being plunge frozen into liquid ethane using a Leica EM GP2 plunge freezer (Leica Microsystems). Frozen grids were imaged using a Titan Krios (Thermo Fisher) operating at 300 keV equipped with a Gatan K3 direct electron detector operating in counting mode.

For the MU89 and MU89+S27Y complexes 1,787 and 1,801 movies, respectively, were collected at a magnification of 81,000x with a physical pixel size of 1.08 Å/pixel using a nominal defocus range of -0.7 to -2.5 µm. Each movie (60 frames) was acquired using a dose rate of ~15.1 $e^-$/pixel/s and a total exposure of ~59 $e^-$/Å2. Motion correction and dose weighting were performed using Unblur(*69*). All subsequent data processing steps including CTF estimation, particle picking, 2D classification, *ab initio* reconstruction, 3D classification and refinement, map local resolution determination were performed in cryoSparc v3 (*70*). Following *ab initio* reconstruction and classification using C1 symmetry, a 3D class was identified with 3 antibody Fabs bound symmetrically to the HIV-1 Env trimer. This initial model was refined using C3 symmetry against the clean stack of particles. Overall map resolution were reported according to the $FSC_{0.143}$ gold-standard criterion (*71*).

*Cryo-EM model fitting***.** Fits of HIV-1 trimer and Fab to the cryo-EM reconstructed maps were performed using ChimeraX (*72*). An initial model for the MU89 Fv region was built using the Abodybuilder-ML website (http://opig.stats.ox.ac.uk/webapps/newsabdab/sabpred/abodybuilder/) (*73*). The Env coordinates from the DH270.6 -bound cryo-EM structure (PDB ID: 6UM6) were used as initial model for Env. To generate coordinates for the MU89+S27Y Fv, the S27Y mutation was introduced into the MU89 coordinates using Coot (*74*). The coordinates were further fit to the electron density first using Isolde (*75*), followed by an iterative process of manual fitting using Coot and real space refinement within Phenix (*76*)**.** Molprobity (*77*) and EMRinger (*78*) were used to check geometry and evaluate structures at each iteration step. Figures were generated in UCSF ChimeraX and PyMOL (The PyMOL Molecular Graphics System, Version 2.0 Schrödinger, LLC).

High-throughput mouse B cell receptor (BCR) repertoire sequencing

Bulk single chain BCR repertoire sequencing of mouse splenocytes was performed as previously described (*59*). Briefly, next-generation sequencing (NGS) was performed on mouse antibody heavy and light chain variable genes using the Illumina MiSeq platform. Total RNA was purified from splenocytes using a RNeasy Mini Kit (Qiagen, Cat# 74104). Purified RNA was quantified via QuBit Flurometer (Thermo Fisher Scientific) and used to generate Illumina-ready heavy and light chain sequencing libraries using the SMARTer Mouse BCR IgG H/K/L Profiling Kit (Takara, Cat# 634422). Briefly, 1 μg of total purified RNA from splenocytes was used for reverse transcription with Poly dT provided in the SMARTer Mouse BCR kit for cDNA synthesis. Heavy and light chain genes were then separately amplified using a 5' RACE approach with reverse primers that anneal in the mouse IgG constant region for heavy chain genes and IgK for the light chain genes (SMARTer Mouse BCR IgG H/K/L Profiling Kit). While the DH270 UCA uses a lambda light chain (VL2-23), the UCA KI mouse model has the light chain gene knocked into the kappa locus, therefore kappa primers provided in the SMARTer Mouse BCR kit were used for light chain gene library preparation for both the DH270 UCA KI and CH235 UCA KI mice. 5 µl of cDNA was used for heavy and light chain gene amplification via two rounds of PCR; PCR1 used 18 cycles and PCR2 used 12 cycles. During PCR2, Illumina adapters and indexes were added. Illumina-ready sequencing libraries were then purified and size-selected by AMPure XP (Beckman Coulter, Cat# A63881) using kit recommendations. The heavy and light chain libraries per mouse were indexed separately, thus allowing us to deconvolute the mouse-specific sequences during analysis. Libraries were validated on the 2200 TapeStation (Agilent) using a D1000 HS kit (Cat# 5067-5584) and quantified a QuBit Fluorometer (Thermo Fisher). Libraries from mice were pooled by groups for sequencing on the Illumina MiSeq Reagent Kit v3 (600 cycle) (Illumina, Cat# MS-102-3003) using read lengths of 301/301 with 20% PhiX.

<u>Antibody sequence analysis</u>

Analysis of the probability of mutations in the absence of antigenic selection was performed using the computational program ARMADiLLO as previously described (*4*). To enable antibody mutation probability analysis of large NGS sequencing datasets, we precomputed per-position amino acid probabilities with ARMADiLLO using the input DH270 UCA sequence and simulated SHM over all possible numbers of mutations (from 1 to N mutations where N represents the sequence length). To analyze each individual antibody sequence, we used lookup tables of the precomputed data which resulted in substantial speed-ups and allowed us to estimate probabilities of mutations observed in all DH270 UCA derived reads recovered from each immunized mouse. Due to variable recovery rates of heavy and light chain reads using the bulk single chain NGS sequencing of mouse BCR repertoires and the depth required for sufficient statistics for analysis of rare mutational events, mouse repertoires that had <1000 unique and functional bnAb UCA knock-in derived reads recovered were excluded from the data analysis (see **Data S2** for number of total and functional bnAb UCA knock-in derived reads recovered for each mouse used in this study). Functional heavy and light chains were defined by the presence of required immunogenetic characteristics as determined by Clonanalyst including: V gene regions of at least 200 base pairs in length, presence of invariant cysteines and tryptophan (heavy) or phenylalanine (light), CDR3s in reading frame 1, non-zero CDR3 length, and absence of stop codons. The use of IgG specific primers ensured IgG isotype recovery for heavy chains, however due to the lack of correspondence between heavy and light chains in bulk single chain BCR repertoire sequencing, we were not able to recover isotype information for light chain repertoire sequencing. As such, mutation frequencies in light chains repertoires are reduced relative to mutation frequencies in the heavy chain because the light chain repertoires include IgM isotypes which include a substantial proportion of unmutated BCRs. Trees of DH270 UCA derived reads representing individual immunized mouse repertoire diversity were generated using 1000 randomly sampled functional heavy chain NGS repertoire reads and were constructed by neighbor-joining using Geneious Prime version 2022.2.1 (https://www.geneious.com) and visualized using the ggtree package in R. In the DH270 clonal tree, antibody intermediates are numbered descending from the number of total ancestral antibody intermediates in the clone. We have previously described a DH270 clonal tree reconstructed from 5 mature members of the DH270 clone (*5*) and after isolation of an additional clonal member, a DH270 clonal tree reconstructed from the 6 member DH270 clone (*36*). Intermediate antibody names are numbered in descending order from the root of the clonal tree and a decimal notation was adopted to distinguish the 6 member DH270 clonal tree from an early reconstruction of the clonal tree that contained 5 members (*5*) e.g. the first antibody intermediate of the six member DH270 clonal tree is named I5.6.

Antibody binding kinetics measurements

*Surface plasmon resonance (SPR).* SPR experiments were performed on a BIACore T200. To determine apparent affinities, approximately 300 RU (range 310-321 RU) of each antibody was captured on an anti-human IgFc immobilized Series S CM5 sensor chip (GE Healthcare). Serial dilutions of SOSIP Env was flowed over immobilized antibody in HEPES buffered saline. To determine binding affinity, biotinylated SOSIP gp140 was immobilized on streptavidin-coated sensor chips. Serial dilutions of antibody Fab were flowed over the Env. Each concentration of Env was flowed over each immobilized antibody for 120 s and dissociation was measured for 600s. In between injections of each Env concentration, the surface was regenerated by injecting glycine pH2 for 30s. Binding rate constants ($k_a$, $k_d$) were measured following global curve fitting to a Langmuir model. Curve fitting analysis was performed with BiaEvaluation software (GE Healthcare) using a 1:1 Langmuir model, or a heterogenous binding model when appropriate to derive rate ($k_a$, $k_d$) and apparent or true equilibrium dissociation constants ($K_d$).

*Biolayer interferometry (BLI).* Biolayer interferometry was performed as previously described (*79*). BLI experiments were conducted on the Octet Red96e system (Sartorius) at 30°C and an orbital shake speed of 1000rpm. All assays used as diluents 0.22μm filtered PBS buffer supplemented with 0.05% Tween 20 and 0.1% bovine serum albumin (PBS-T-BSA) and flat bottom 96-well plates (Greiner).

For the ligand titration experiment, two-fold serial dilutions of the biotinylated CH848 10.17.SOSIP and CH848 10.17DT.SOSIP trimers, with a starting concentration of 10μg/mL, were immobilized on hydrated Streptavidin (SA) Tips (Octet® Sartorius) for 120 seconds. After a 60 second wash and 180 second Baseline step in PBS-T-BSA, the tips were incubated with 2000nM of DH270 UCA Fab, 500nM of Mu89 Fab and 500nM of Mu89+S27Y Fab for 600 seconds. A ligand-coated sensor tip dipped into buffer only was used as the negative reference. The negative reference values were subtracted from all values obtained from wells that included Fabs. The optimal Env concentrations for affinity studies were determined by analyzing the sensorgram traces and binding responses using Data Analysis HT 12.0 software (Forte Bio). For the binding kinetics experiment, the chosen Env concentrations were immobilized on hydrated SA biosensor tips and incubated with two-fold serial dilutions of antibody Fabs for 600 seconds followed by a 600 second long dissociation step in PBS-T-BSA. A ligand-coated sensor tip dipped into buffer only was used as the negative reference sensor. Background values obtained with the negative reference sensor were subtracted from all values obtained for sensors submerged in wells containing Fabs. The binding response sensorgram curves were globally fitted using a 1:1 Binding Model and the rate constants $k_a$, $k_d$ and $K_d$ were calculated using Data Analysis HT 12.0 software (Forte Bio). In some instances, interst

For antibody binding breadth analyses, 20 μg/mL IgG was immobilized to anti-human Fc (AHC) sensor tips (Octet® Sartorius) and incubated with 50 μg/mL of Env SOSIP gp140 trimers or Env SOSIP gp140-nanoparticles for 400 seconds. Wavelength shift (nm) at the end of association was calculated in Data Analysis HT 12.0 software (Forte Bio) and used to determine positive binding.

SARS-CoV-2 spike was used as a negative analyte and anti-influenza antibody CH65 was used as the negative ligand reference. CH65 values were reference subtracted from all test antibodies. Antibody binding was considered positive when the nm value was greater than 0.02 nm shift and at least three times higher than the SARS-CoV-2 spike values used as the negative analyte.

Statistical analysis

Differences between mouse groups were tested for statistical significance using two-sided Wilcoxon Mann-Whitney exact tests in R or GraphPad Prism and p-values are reported without multiple test corrections.

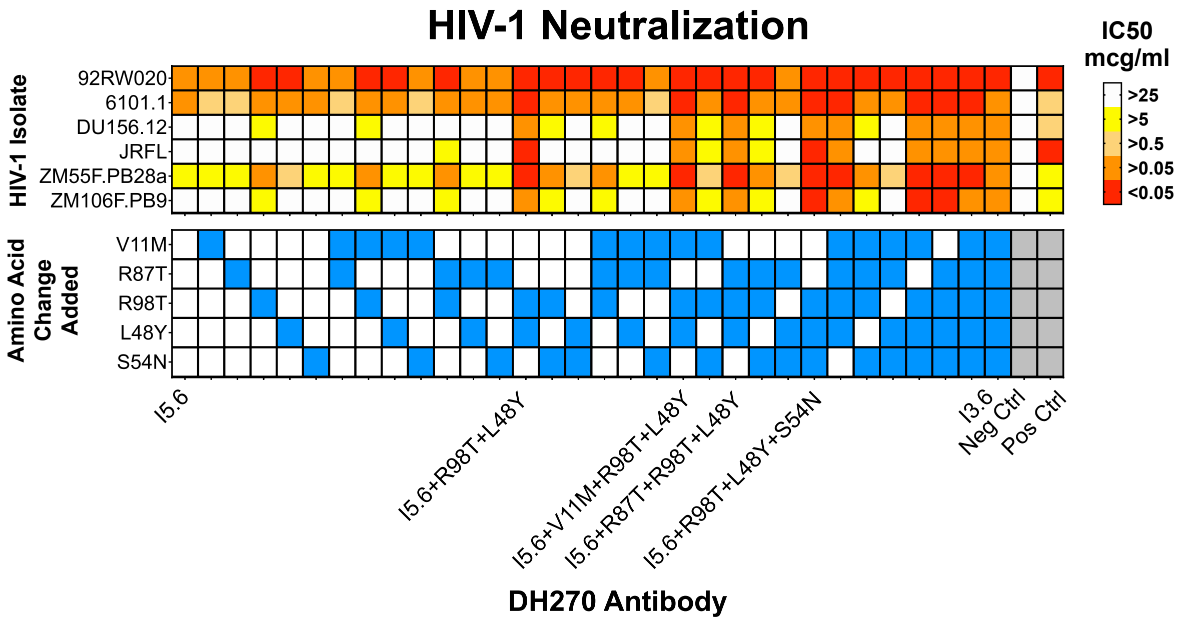


**Fig. S1. Neutralization of antibody mutants with all combinations of 5 mutations from the second intermediate I3.6 introduced into the first intermediate I5.6 of the DH270.6 clonal lineage.** Neutralization of 7 heterologous viruses that are sensitive to I3.6 neutralization (upper panel) with each mutation combination denoted by blue filled boxes (lower panel). Neg ctrl = CH65 antibody; Pos ctrl = CH31 and CH01 antibodies.

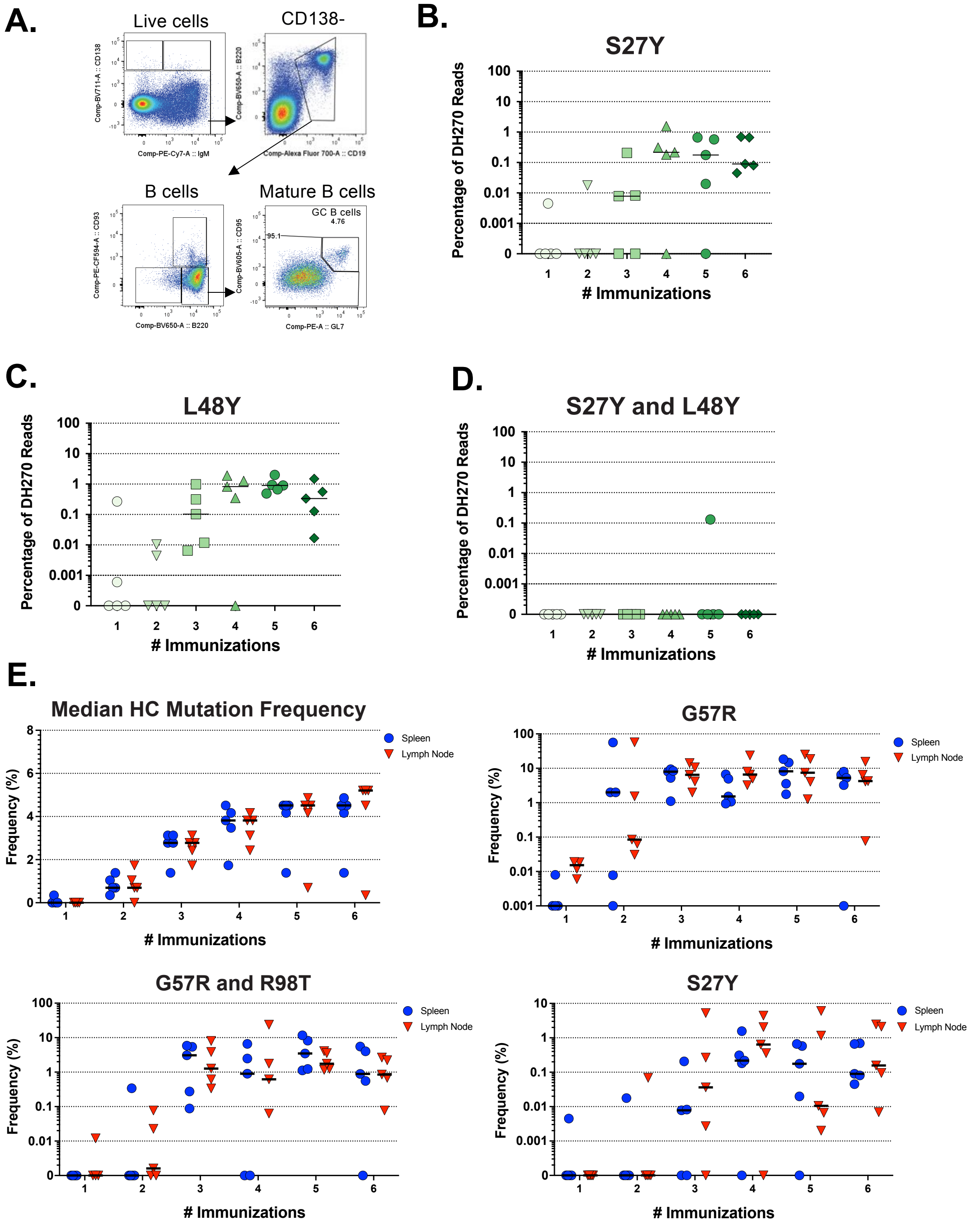
A.
Live cells
CD138-
B cells
Mature B cells
GC B cells
4.76
95.1
Comp-BV711-A :: CD138
Comp-PE-Cy7-A :: IgM
Comp-BV650-A :: B220
Comp-Alexa Fluor 700-A :: CD19
Comp-PE-CF594-A :: CD93
Comp-BV650-A :: B220
Comp-BV605-A :: CD95
Comp-PE-A :: GL7
B.
S27Y
Percentage of DH270 Reads
### Immunizations
C.
L48Y
Percentage of DH270 Reads
### Immunizations
D.
S27Y and L48Y
Percentage of DH270 Reads
### Immunizations
E.
Median HC Mutation Frequency
Frequency (%)
### Immunizations
Spleen
Lymph Node
G57R
Frequency (%)
### Immunizations
Spleen
Lymph Node
G57R and R98T
Frequency (%)
### Immunizations
Spleen
Lymph Node
S27Y
Frequency (%)
### Immunizations
Spleen
Lymph Node

F.

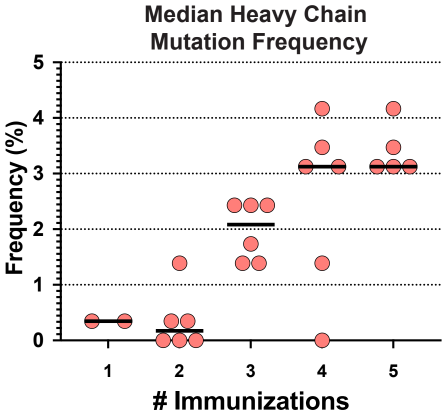


G.

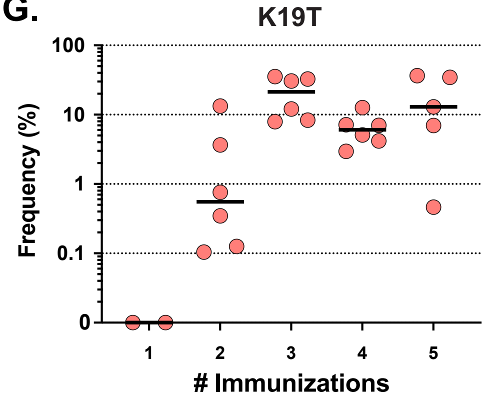


H.

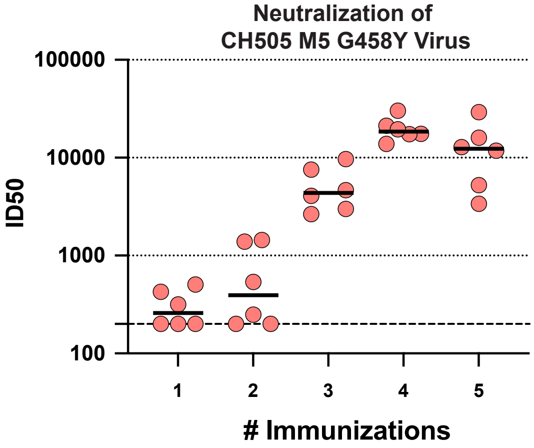


**Fig. S2. Mutation frequencies and neutralization profiles in response to repetitive immunizations.** A) Flow cytometry gating strategy for germinal center B cells in mice immunized 1-6 times with the 10.17 DT nanoparticle immunogen. Frequencies of early-lineage key improbable light chain mutations B) S27Y C) L48Y and D) S27 and L48Y in combination in response to 1-6 immunizations with the 10.17DT nanoparticle. Similar dynamics of mutation acquisition was observed between draining lymph nodes and spleens as measured by BCR repertoire sequencing for E) median heavy chain mutation frequencies and frequencies of key improbable mutations F) Median heavy chain mutation frequency G) K19T improbable mutation frequency and H) neutralization titers to the vaccine-matched CH505 M5 G458Y virus in response to repetitive immunizations of the CH505 M5/G458Y/GNTI- Env SOSIP trimer immunogen in CH235 UCA knockin mice.

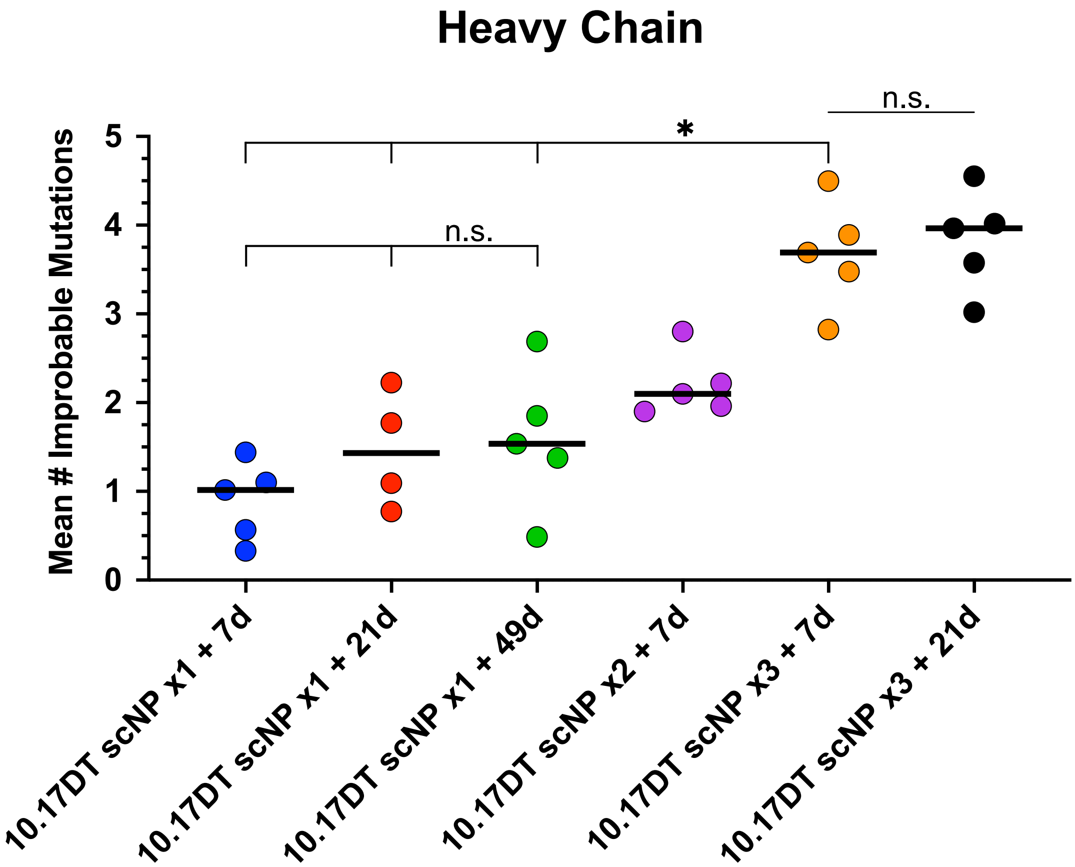


**Fig. S3. Mutation acquisition dynamics in response to single immunizations with varying maturation times vs. repetitive immunization with the 10.17DT nanoparticle.** A) Mean number of improbable mutations acquired by mice immunized with single immunizations of the CH848 10.17DT SOSIP nanoparticle (10.17DT scNP) and BCR repertoire sequenced 7, 21 or 50 days later or mice immunized with 2-3 times and sequenced 7 days later or immunized 3 times and sequenced 21 days later. Each dot represents one mouse. * $p<0.05$; Wilcoxon-Mann-Whitney test. Statistical significance only shown for group comparisons referenced in the main text.

A.

| Antibody | 10.17 DT | | | 10.17 WT | | |
|---|---|---|---|---|---|---|
| | ka (1/Ms) | kd (1/s) | KD (nM) | ka (1/Ms) | kd (1/s) | KD (nM) |
| UCA3* | 4245 | 2.06E-03 | 485.7 | 3500 | 4.89E-03 | 1400 |
| UCA3+G57R | 2477 | 3.94E-04 | 159.2 | 679.7 | 3.63E-04 | 533.4 |
| I5.6* | 4734 | 1.58E-04 | 33.29 | 944.4 | 1.65E-04 | 175 |
| UCA3+G57R+R98T | 3143 | 4.11E-05 | 13.1 | 1046 | 5.45E-05 | 52.12 |
| I3.6* | 2807 | 9.38E-05 | 33.39 | 1451 | 2.36E-04 | 162.8 |
| I2.6* | 12430 | 1.61E-05 | 1.29 | 8141 | 1.05E-05 | 1.29 |
| DH270.6* | 16740 | 1.03E-05 | 0.62 | 11470 | 7.32E-06 | 0.64 |

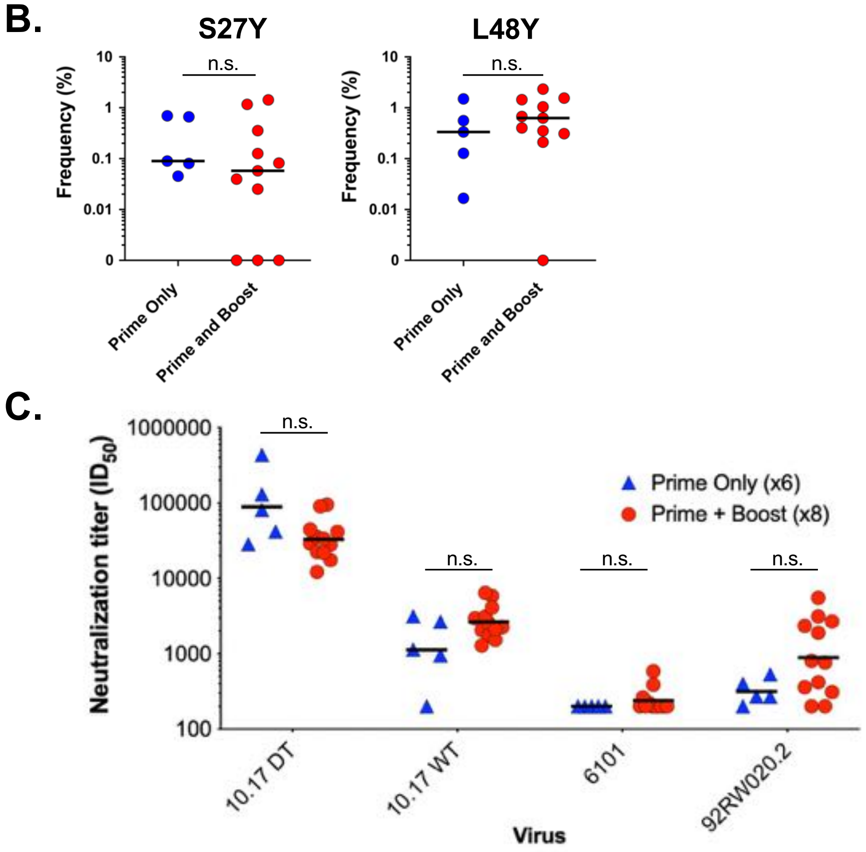


**Fig. S4. Binding kinetics of prime-boost immunogens, light chain mutation frequencies, and neutralization of primed and boosted mice.** A) Binding kinetics measurements of DH270 lineage antibodies and antibody mutants against prime (10.17DT) and boost (10.17 WT) immunogens measured by SPR. *Kinetics data for these antibodies is reported in submitted manuscript https://www.biorxiv.org/content/10.1101/2022.09.14.507935v1 B) Frequency of DH270 reads with key light chain improbable mutations S27Y (left) and L48Y (right) in prime only and prime and boost mice. C) Neutralization of vaccine-matched (priming immunogen) 10.17 DT, V1 glycans-containing 10.17 WT (vaccine-matched as boosting immunogen in prime-boost regimen group), and heterologous tier 1B viruses 6101 and 92RW020. Each dot represents one immunized mouse in panels B and C.

#### Heavy Chain

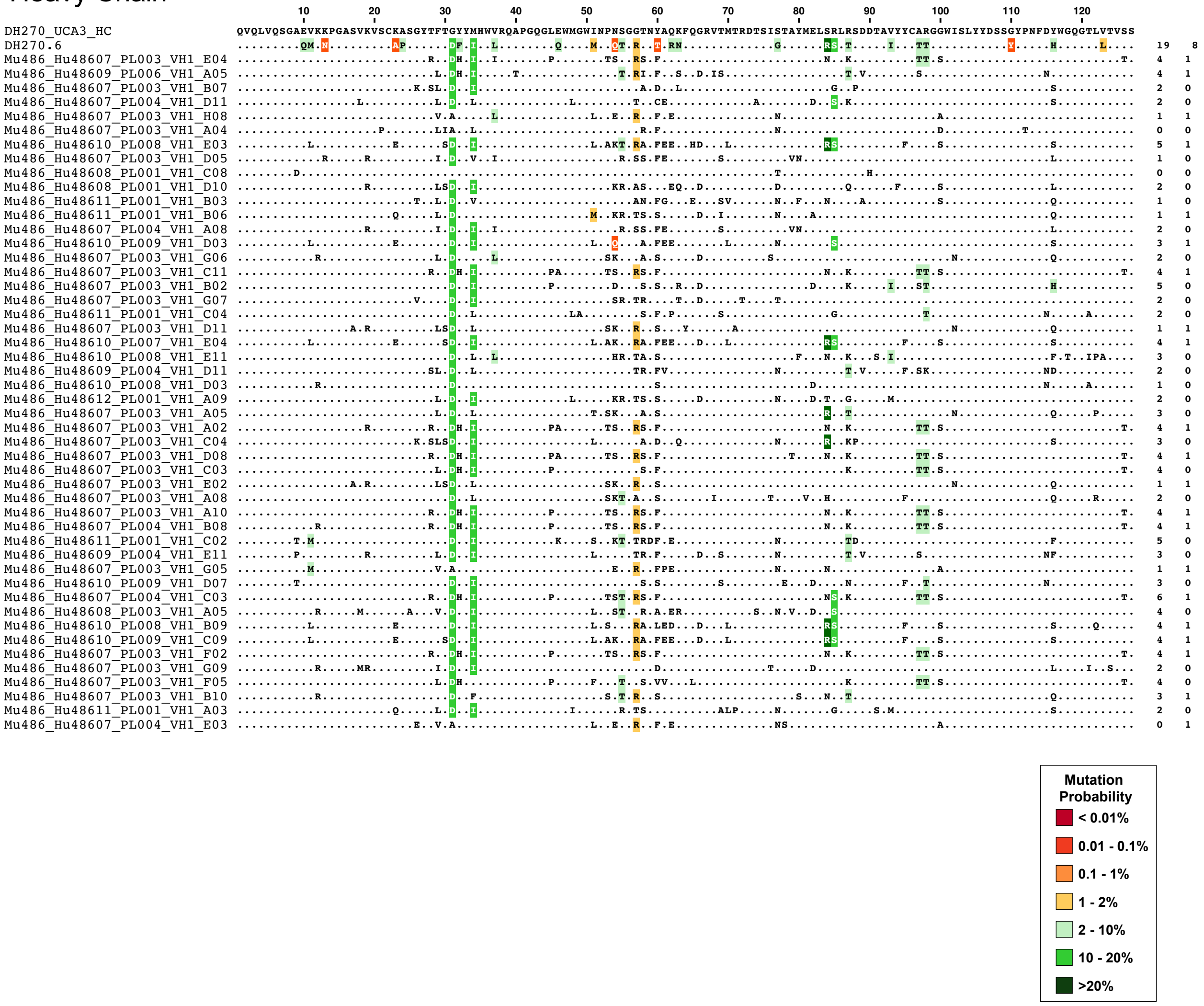
DH270_UCA3_HC
QVQLVQSGAEVKKPGASVKVSCKASGYTFTGYYMHWVRQAPGQGLEWMGWINPNSGGTNYAQKFQGRVTMTRDTSISTAYMELSRLRSDDTAVYYCARGGWISLYYDSSGYPNFDYWGQGTLVTVSS
DH270.6
Mu486_Hu48607_PL003_VH1_E04
Mu486_Hu48609_PL006_VH1_A05
Mu486_Hu48607_PL003_VH1_B07
Mu486_Hu48607_PL004_VH1_D11
Mu486_Hu48607_PL003_VH1_H08
Mu486_Hu48607_PL003_VH1_A04
Mu486_Hu48610_PL008_VH1_E03
Mu486_Hu48607_PL003_VH1_D05
Mu486_Hu48608_PL001_VH1_C08
Mu486_Hu48608_PL001_VH1_D10
Mu486_Hu48611_PL001_VH1_B03
Mu486_Hu48611_PL001_VH1_B06
Mu486_Hu48607_PL004_VH1_A08
Mu486_Hu48610_PL009_VH1_D03
Mu486_Hu48607_PL003_VH1_G06
Mu486_Hu48607_PL003_VH1_C11
Mu486_Hu48607_PL003_VH1_B02
Mu486_Hu48607_PL003_VH1_G07
Mu486_Hu48611_PL001_VH1_C04
Mu486_Hu48607_PL003_VH1_D11
Mu486_Hu48610_PL007_VH1_E04
Mu486_Hu48610_PL008_VH1_E11
Mu486_Hu48609_PL004_VH1_D11
Mu486_Hu48610_PL008_VH1_D03
Mu486_Hu48612_PL001_VH1_A09
Mu486_Hu48607_PL003_VH1_A05
Mu486_Hu48607_PL003_VH1_A02
Mu486_Hu48607_PL003_VH1_C04
Mu486_Hu48607_PL003_VH1_D08
Mu486_Hu48607_PL003_VH1_C03
Mu486_Hu48607_PL003_VH1_E02
Mu486_Hu48607_PL003_VH1_A08
Mu486_Hu48607_PL003_VH1_A10
Mu486_Hu48607_PL004_VH1_B08
Mu486_Hu48611_PL001_VH1_C02
Mu486_Hu48609_PL004_VH1_E11
Mu486_Hu48607_PL003_VH1_G05
Mu486_Hu48610_PL009_VH1_D07
Mu486_Hu48607_PL004_VH1_C03
Mu486_Hu48608_PL003_VH1_A05
Mu486_Hu48610_PL008_VH1_B09
Mu486_Hu48610_PL009_VH1_C09
Mu486_Hu48607_PL003_VH1_F02
Mu486_Hu48607_PL003_VH1_G09
Mu486_Hu48607_PL003_VH1_F05
Mu486_Hu48607_PL003_VH1_B10
Mu486_Hu48611_PL001_VH1_A03
Mu486_Hu48607_PL004_VH1_E03
Mutation Probability
< 0.01%
0.01 - 0.1%
0.1 - 1%
1 - 2%
2 - 10%
10 - 20%
>20%

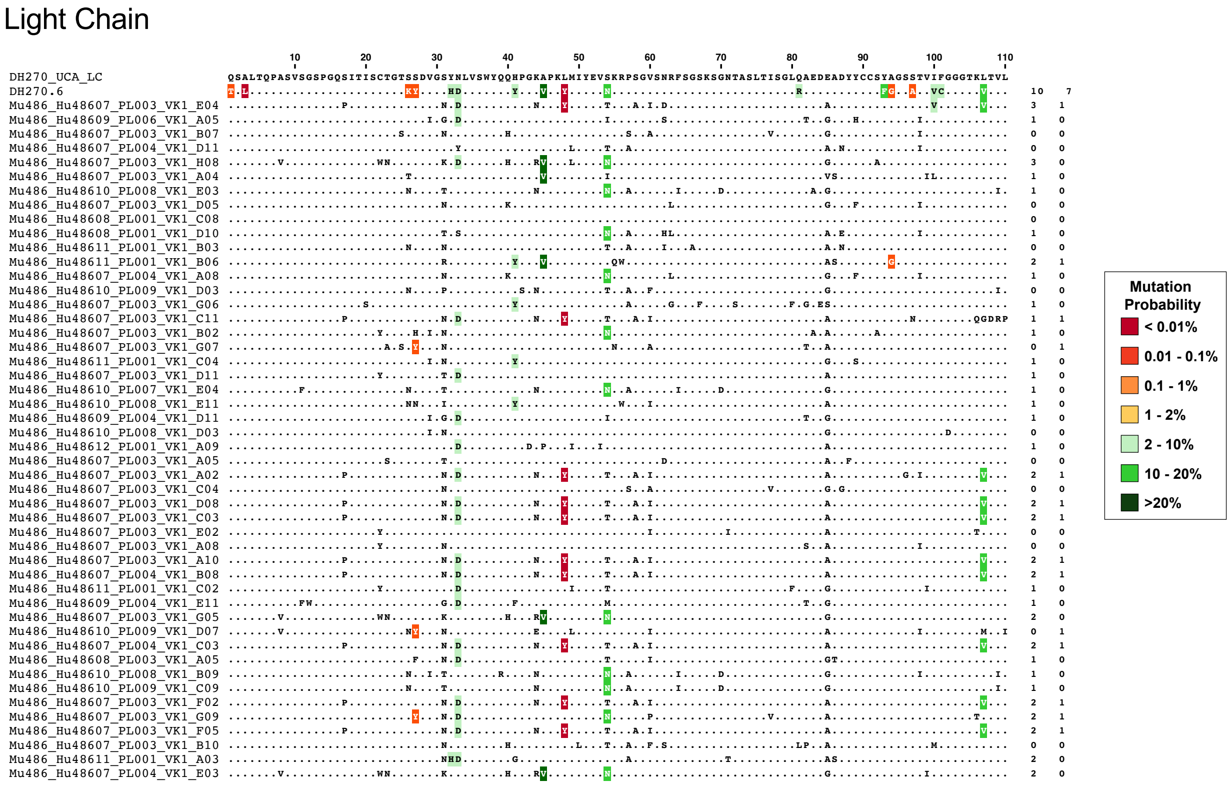


**Fig. S5. Prime-boost immunization elicited antibodies with somatic mutations shared with DH270.6.** Heavy chain (upper panel) and light chain (lower panel) sequences of isolated DH270 UCA knockin derived monoclonal antibodies aligned to DH270 UCA and DH270.6. Dots represent amino acids that are in common with DH270 UCA. Mutations shared with DH270.6 are highlighted in colors according to their mutation probability as estimated by ARMADiLLO. The numbers to the right of each row are total numbers of probable mutations and improbable mutations observed in the sequence, respectively.

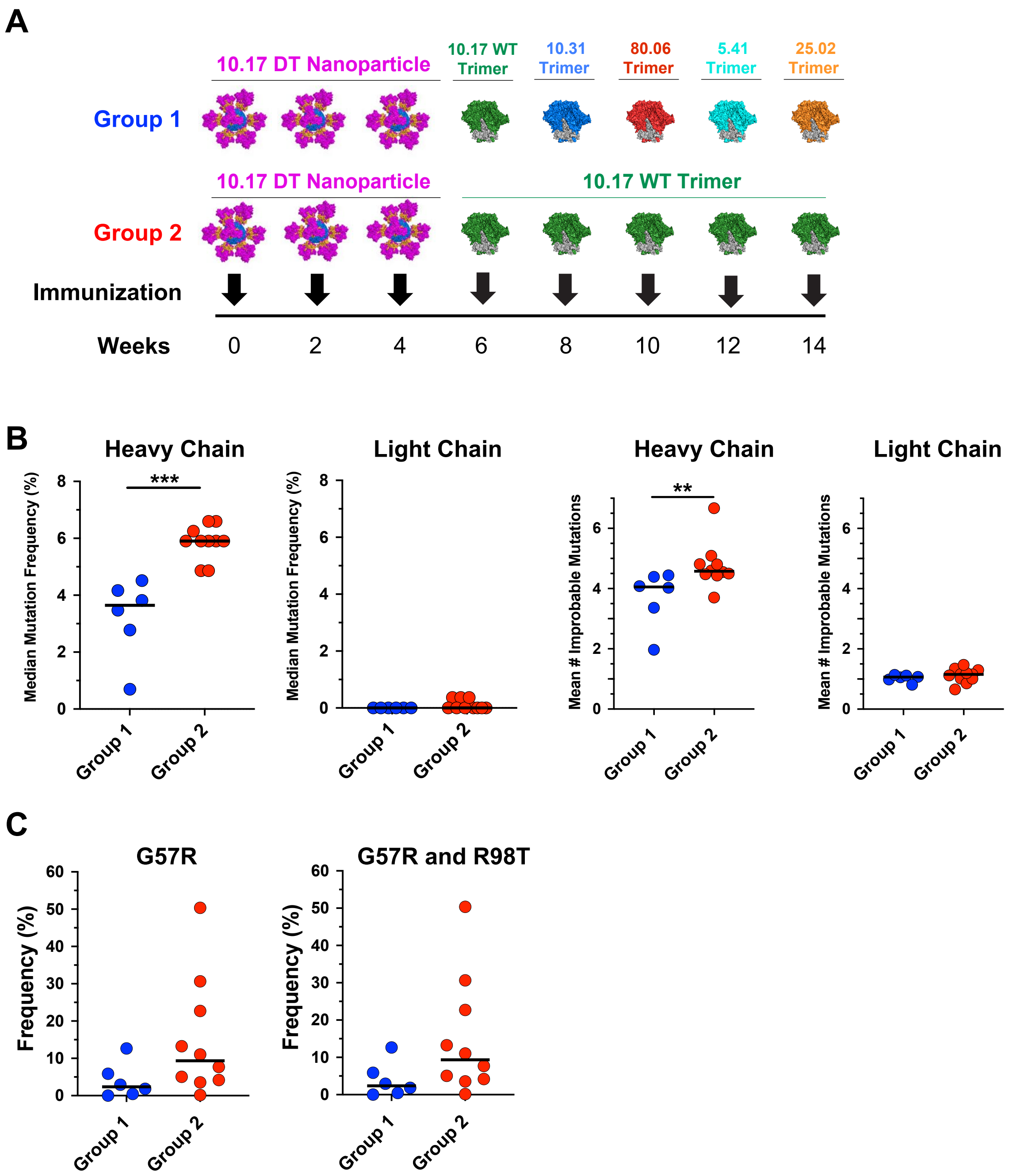


**Fig. S6. Comparison of frequencies of mutations selected by prime-boost regimens.** A) Schematic of immunization schedules for prime-boost regimens. Mouse groups 1 and 2 were primed 3 times with the 10.17DT nanoparticle. Group 1 mice were boosted with a single 10.17WT SOSIP trimer boost followed by single boosts with four autologous CH848 virus Env SOSIP trimers each. Group 2 was boosted 5 times with the 10.17WT SOSIP trimer B) median mutation frequency and mean number of improbable mutation for heavy and light chains as measured by mouse BCR repertoire sequencing for immunized mice C) frequency of key heavy chain improbable mutations. *** p-value <0.001, Wilcoxon-Mann-Whitney exact test.

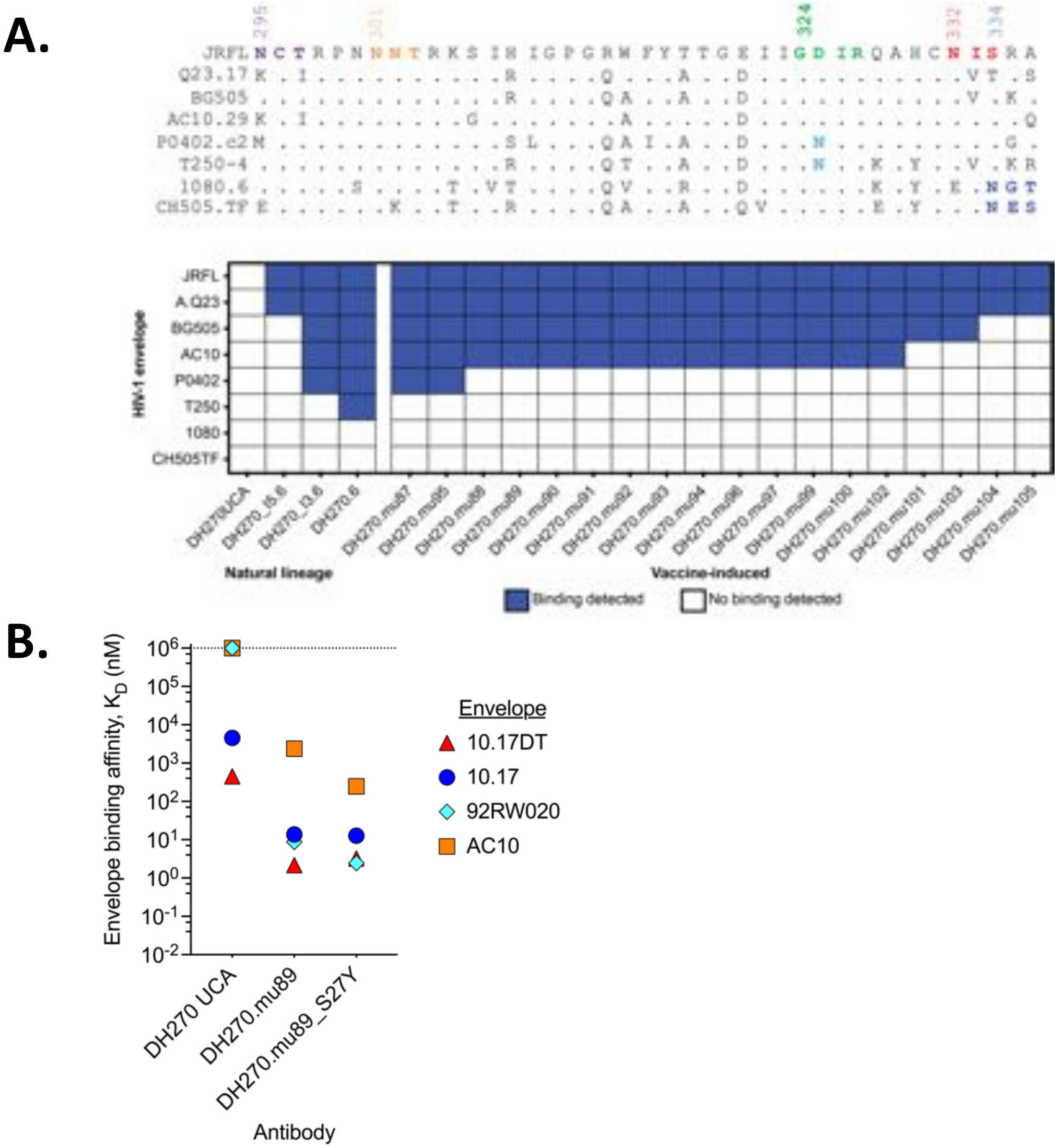


**Fig. S7. Affinity Maturation of Monoclonal antibodies elicited by 10.17DT nanoparticle prime and 10.17WT SOSIP trimer boost regimen results in Broad Env Reactivity and high affinity for Heterologous HIV-1 envelopes.** A) Vaccine-induced DH270 antibody binding breadth determined for a multi-clade panel of stabilized gp140 envelopes. Envelope V3 glycan site sequence alignment is shown at the top. Identical amino acids are shown as periods. Glycosylation sites are highlighted in various colors. The GDIR motif is shown in green. Positive or negative antibody binding measured by biolayer interferometry is shown below the sequence alignment. Binding breadth of sequential members of the natural DH270 lineage ordered from least to most somatically mutated is shown in the first four columns of the graph to demonstrate the acquisition of binding breadth concurrent with further somatic mutation. B) Binding affinity shown as equilibrium dissociation constant ($K_D$) of DH270.mu89 with and without the S27Y substitution for autologous (10.17DT and 10.17) and heterologous (92RW020 and AC10)

stabilized gp140 envelopes. DH270.mu89 has affinity matured to bind heterologous envelopes, which can be improved by the addition of the S27Y substitution. The affinities of DH270 UCA for each envelope are shown to indicate the DH270 lineage initial affinities for each envelope.

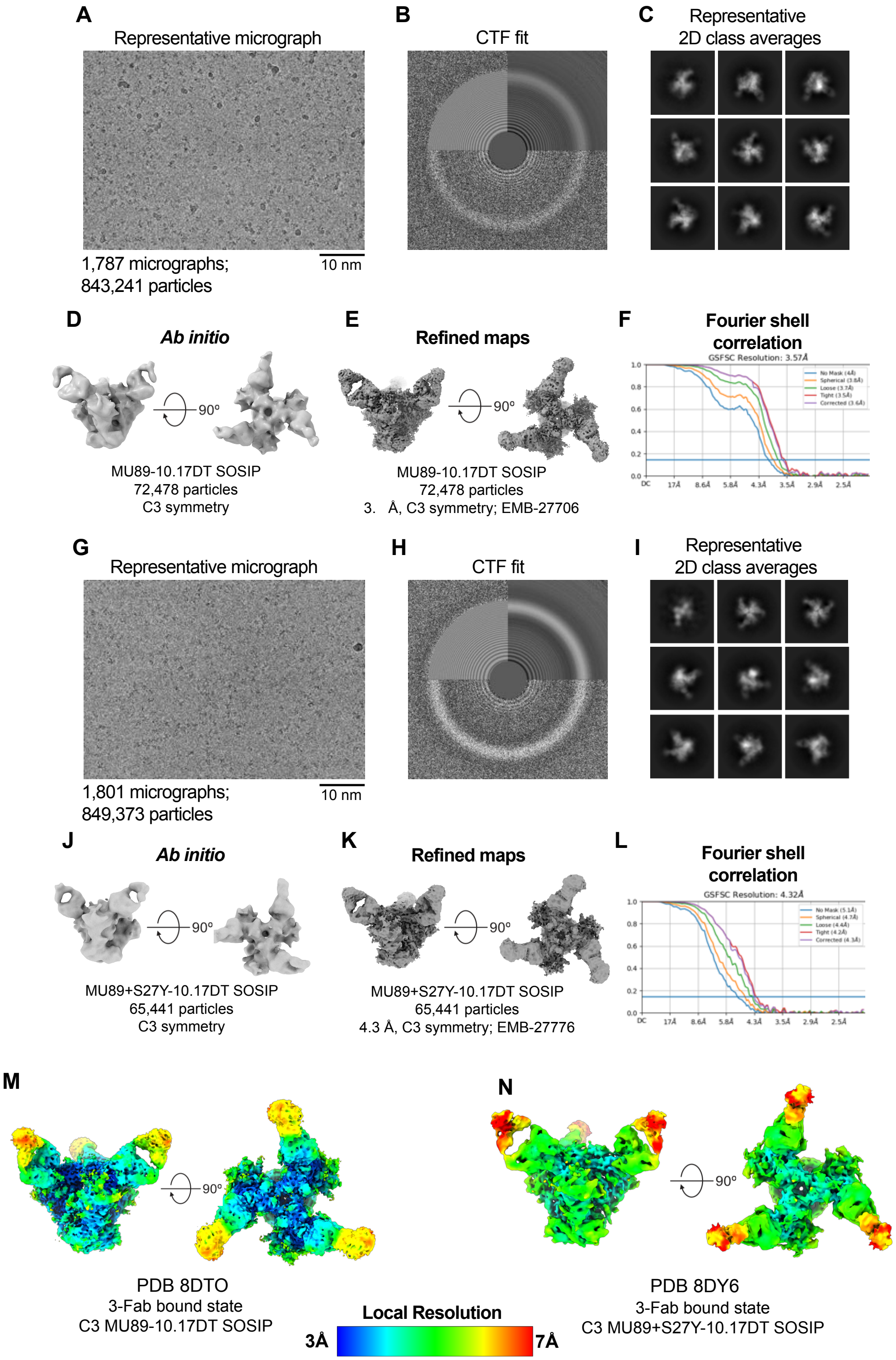


**Fig. S8. Cryo-EM data processing for the MU89 and MU89+S27Y bound complexes to HIV-1 Env Trimer.** (A-F) MU89 bound to 10.17DT SOSIP (A) Representative micrograph. (B)

CTF Fit. (C) Representative 2D class averages from cryo-EM dataset. Box size = 345.6 Å. (D) Ab initio reconstructions for the cryo-EM 3-Fab bound state. (E) Refined maps for the cryo-EM corresponding state. (F) Fourier shell correlation curves for the cryo-EM corresponding state. (G-L) MU89+S27Y bound to 10.17DT SOSIP (G) Representative micrograph. (H) CTF Fit. (I) Representative 2D class averages from cryo-EM dataset. Box size = 345.6 Å. (J) Ab initio reconstructions for the cryo-EM 3-Fab bound state. (K) Refined maps for the cryo-EM corresponding state. (L) Fourier shell correlation curves for the cryo-EM corresponding state. (M) Refined maps colored by local resolution ranging from 3 Å to 7 Å.

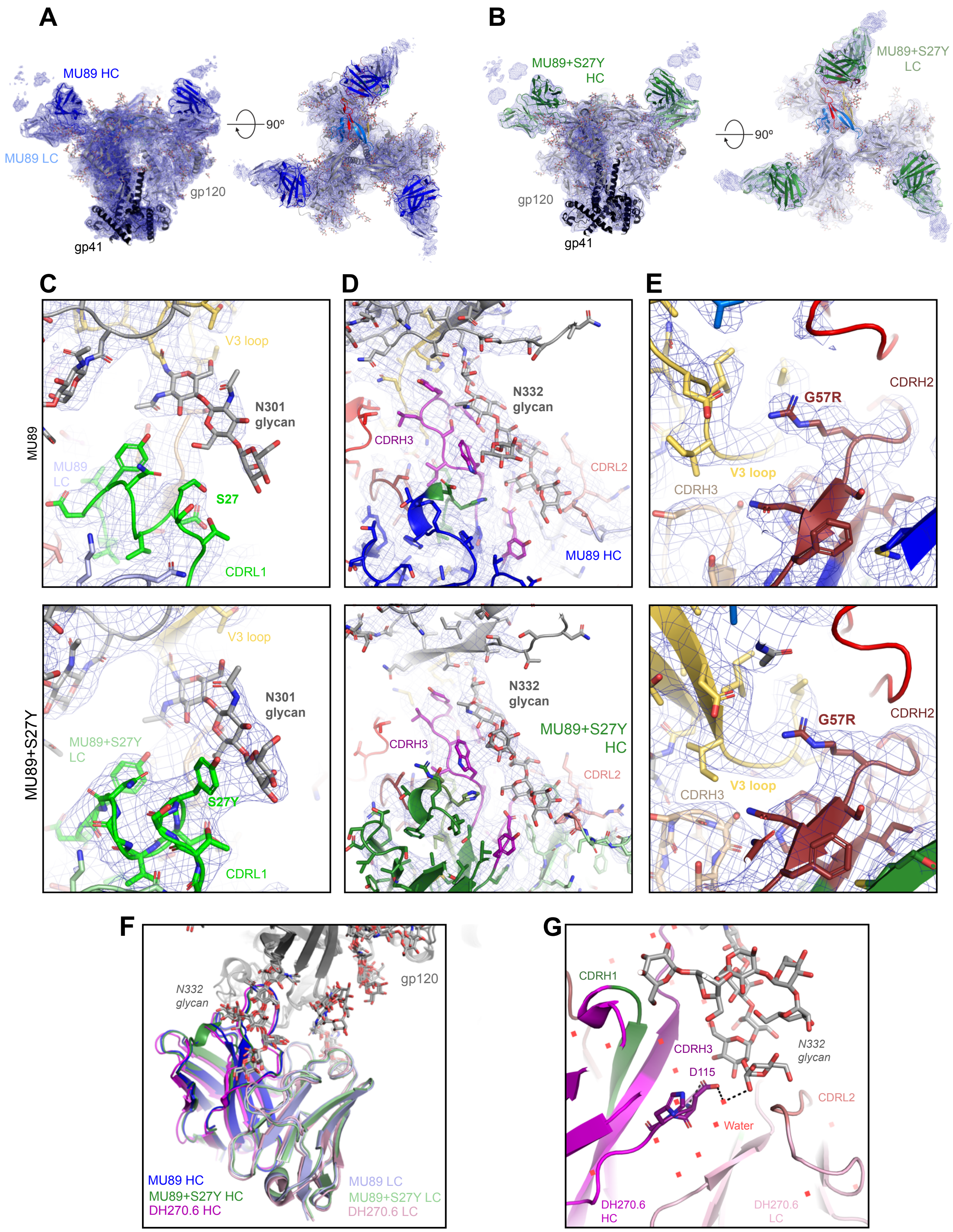
A
MU89 HC
MU89 LC
gp120
gp41
90º
B
MU89+S27Y HC
MU89+S27Y LC
gp120
gp41
90º
C
D
E
MU89
MU89+S27Y
V3 loop
N301 glycan
MU89 LC
S27
CDRL1
N332 glycan
CDRH3
CDRL2
MU89 HC
G57R
CDRH2
V3 loop
CDRH3
V3 loop
N301 glycan
MU89+S27Y LC
S27Y
CDRL1
N332 glycan
MU89+S27Y HC
CDRH3
CDRL2
G57R
CDRH2
CDRH3
V3 loop
F
N332 glycan
gp120
MU89 HC
MU89+S27Y HC
DH270.6 HC
MU89 LC
MU89+S27Y LC
DH270.6 LC
G
CDRH1
CDRH3
D115
N332 glycan
CDRL2
Water
DH270.6 HC
DH270.6 LC

**Fig. S9. Structural details of Env binding interfaces of vaccine elicited antibodies MU89 and MU89+S27Y.** A. Side and top views of MU89 Fab bound to 10.17 DT SOSIP Env trimer cryo-EM model with electron density map shown in blue mesh. MU89 HC (blue), MU89 LC (light blue), gp120 (gray), and gp41 (black). PDB ID: 8DTO, EMD-27706. B. Side and top views of MU89+S27Y Fab bound to 10.17 DT SOSIP Env trimer cryo-EM model with electron density map shown in blue mesh. MU89+S27Y HC (dark green) and LC (pale green). PDB ID: 8DY6, EMD-27776. C. Zoomed-in views of $V_L$ S27Y interaction with N301 glycan. Top MU89 with S27 and bottom MU89+S27Y with S27Y. D. Fab interactions with N332 glycan. E. Zoomed-in views of $V_H$ G57R interaction with the V3 loop. F. Antibody binding overlay aligned by gp120 chains. MU89 (PDB ID: 8DTO), MU89+S27Y (PDB ID: 8DY6), and DH270.6 (PDBID: 6UM6). G. DH270.6 crystal structure (PDB ID:6CBP) highlighting a water mediated interaction of $V_H$ D115 in the HCDR3 region CDRH3 with the N332 glycan.

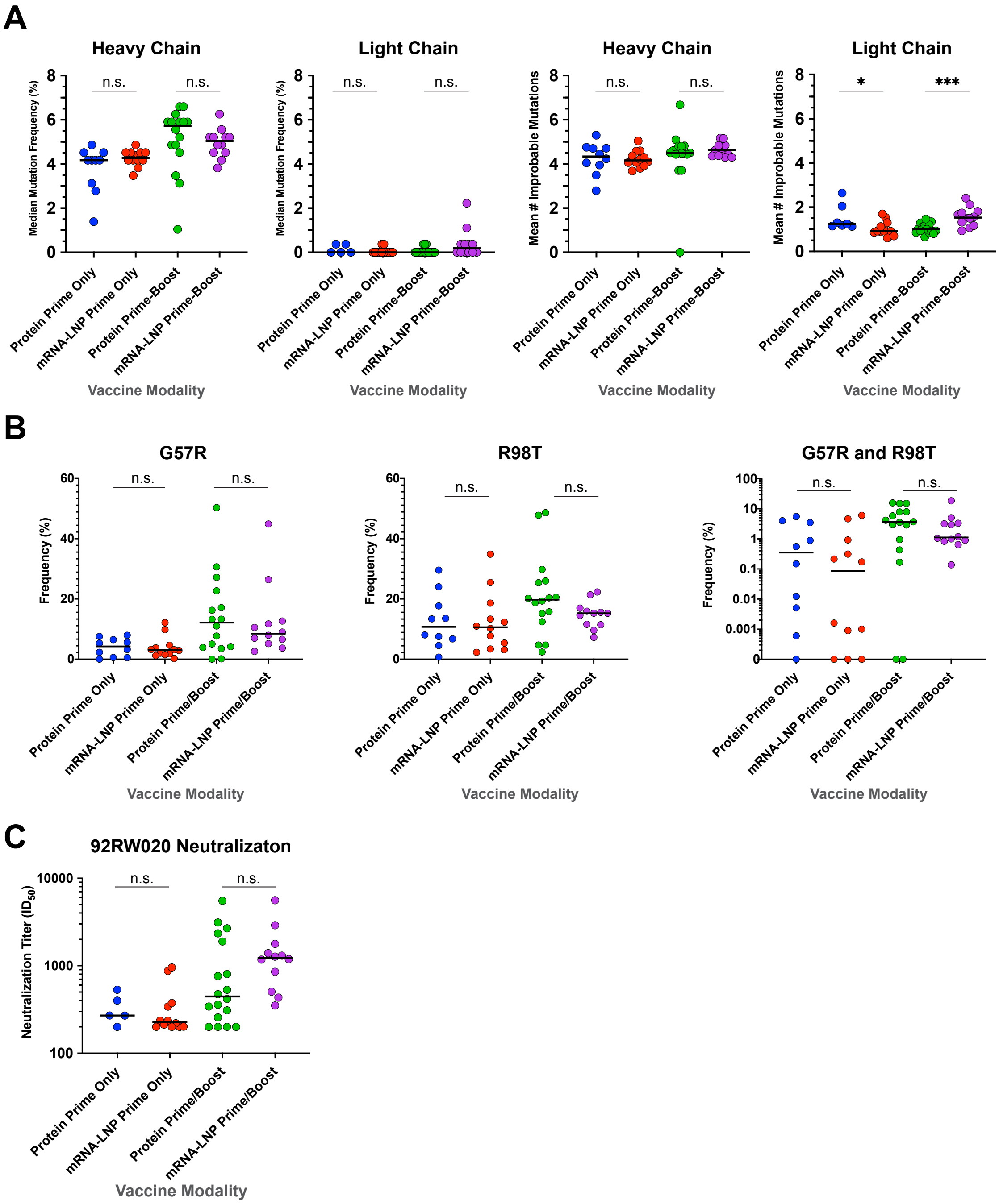


**Fig. S10. Highly similar mutational responses in mRNA-LNP and protein immunized mouse groups for both prime-only and prime-boost regimens.** A) Median mutation frequency (computed as nucleotide mutations in VH or VL gene segment) of DH270 heavy and light chain reads (left two panels) and mean number of heavy and light chain improbable amino acid mutations (right two panels) for pooled groups of mice that received protein prime-only, protein prime-boost, mRNA prime-only and mRNA prime-boost immunization regimens. B) Frequency in the repertoire of DH270 UCA derived reads containing key improbable heavy chain mutations G57R, R98T and their combination. C) Heterologous virus (92RW020) neutralization. Data for the protein prime only group and mRNA prime only group were originally reported in Saunders et al. Science 2019, and Mu et al. Cell Reports 2022, respectively. See Data S1 for details of regimens used in the pooled mouse groups. Each dot represents one mouse. For visual clarity, bars reporting statistical significance testing are only shown for pairwise group comparisons referenced in the main text. See Data S3 for p values of all pairwise group comparisons. **** $p<0.0001$; Wilcoxon-Mann-Whitney test.

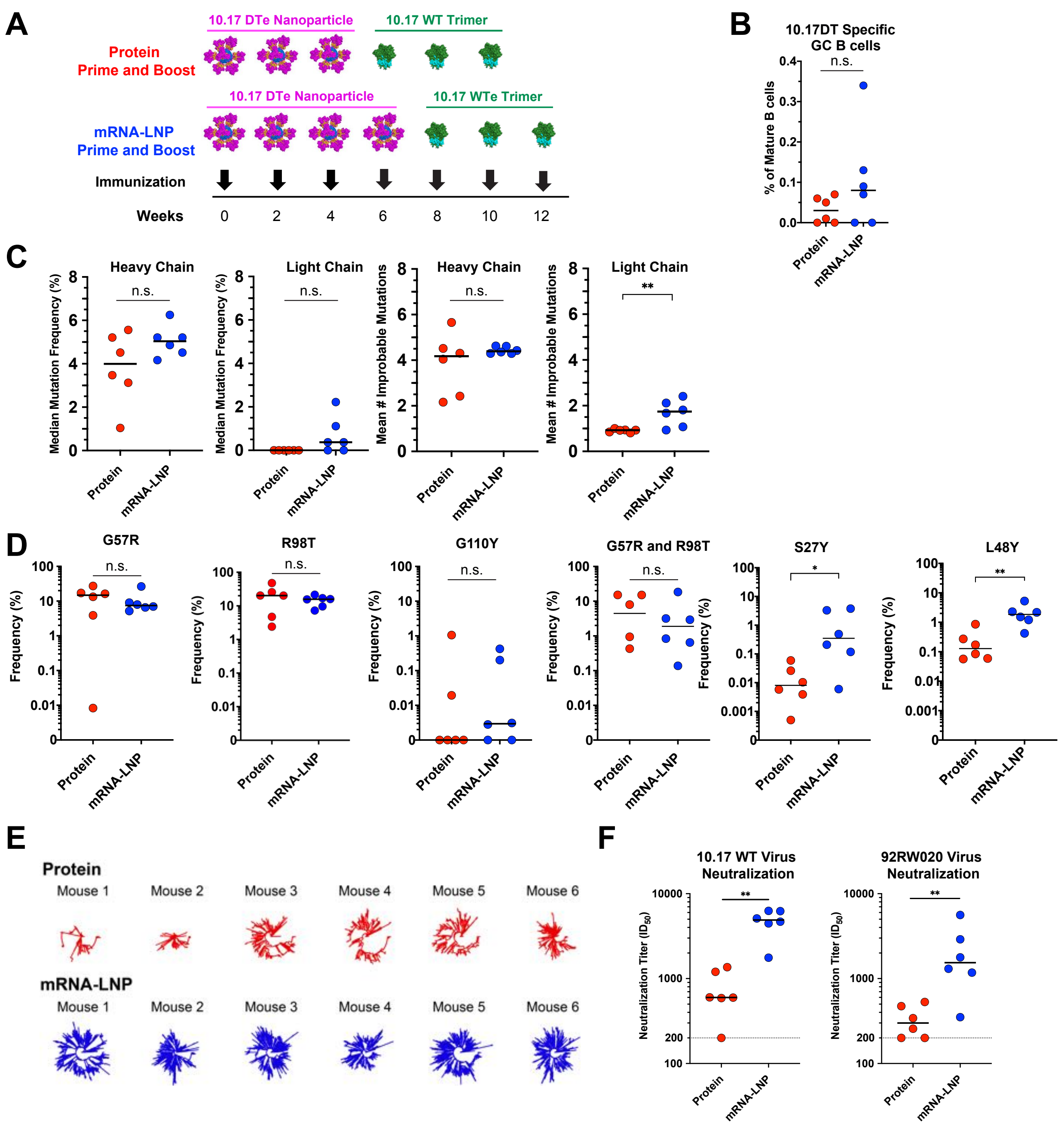


**Fig. S11. mRNA-LNP prime-boost regimen results in higher frequency of light chain improbable mutations and higher heterologous neutralization than protein prime-boost regimen.** A) Schematic of mRNA-LNP and protein prime-boost protein regimens (n=6 mice per group). B) Percentage of GC B cells in the mature B cell population as measured by flow cytometry in protein prime-boost vs. mRNA-LNP prime-boost immunized mice. C) Median mutation frequency (computed as nucleotide mutations in VH or VL gene segment) of DH270 heavy and light chain reads (left two panels) and mean number of heavy and light chain improbable amino acid mutations (right two panels). D) Frequency in the repertoire of DH270 UCA derived reads containing individual key improbable mutations. E) DH270 clonal diversification as shown by circular trees of randomly sampled DH270 reads from the repertoires of each mouse in the mRNA and protein immunization groups. F) Vaccine boost-matched virus (10.17WT) and heterologous virus (92RW020) neutralization. * p<0.05; ** p<0.01; Wilcoxon-Mann-Whitney test.

**Table S1. Cryo-EM data collection and refinement statistics**

**Data S1. (separate file)**
Type or paste caption here.

**Data S2. (separate file)**

**Data S3. (separate file)**
